# Variabilities in the shapes of the receptive fields of simple cells in mice

**DOI:** 10.64898/2026.10.07.757489

**Authors:** Daria Meshcherina, Benjamin Auffarth, Tony Lindeberg

## Abstract

Observing an object under different viewing conditions, in spite of substantial variability in the image structures when changing the distance or the viewing direction relative to the object, we can maintain the identity of the object. How does the vision system cope with this variability, that will substantially influence the receptive field responses? One hypothesis is that the receptive field family should be covariant under geometric image transformations and thereby have variabilities in the shapes of the receptive fields corresponding to the degrees of freedom of the geometric image transformations. Specifically, it has been predicted that the receptive fields should have a variability in the degree of elongation, corresponding to varying the slant angle when observing local surface patches. Such a variability has also been predicted from indirect support in terms of variabilities in the orientation selectivity properties in V1 for monkeys and cats, which by a theoretical model has been linked to variabilities in the elongation of the receptive fields. In this paper, we measure such variabilities in receptive field shapes in a more direct way, by reconstructing receptive fields from calcium imaging experiments for mice previously performed by the Allen Institute. Our findings reveal a variability in the degree of elongation that exceeds what measurement noise produces, spanning about a factor of 5, thereby showing that the predictions that were made for higher mammals with orientation maps, such as primates and cats, also apply to a species without orientation maps, with its primary visual cortex instead based on a salt-and-pepper organization of the receptive fields.

**Author summary:** The primary visual cortex contains cells that each respond to a small patch of the visual scene. For the visual system to recognise that two different views of the same object belong together, these cells must come in a range of shapes that between them cover the ways an image can change as we, or the objects around us, move. A mathematical theory of vision predicts which shapes should be present, and in particular that the cells should vary in how elongated they are. That prediction had only been tested indirectly, in animals such as cats and monkeys whose visual cortex is arranged into orderly maps of shape preference. We tested for this hypothesis by more direct means for the mouse, whose visual cortex has no such map, by fitting a mathematical model to visual cortex neurons recorded in a large public dataset. Among the cells we could confirm as the simplest type, the degree of elongation varied by more than measurement noise alone can produce, spanning roughly a factor of five, scattered across the cortex with no orderly arrangement.

## Introduction

How a visual system copes with the geometry of viewing has been approached from two broad directions. One places the problem downstream of the primary visual cortex: Marr [1] treated vision as a staged reconstruction, from a primal sketch through a viewer-centred description of surfaces to an object-centred model. The division of labour between a ventral and a dorsal stream (Ungerleider and Mishkin [2], Goodale and Milner [3]) locates the resolution of viewpoint well beyond the front end, as does more recent work that recovers scene parameters by inverting a generative model (Yildirim *et al*. [4]); and in artificial networks (Zhu *et al*. [5]). The other derives the shapes of early receptive fields from the statistics of natural images, through redundancy reduction (Barlow [6]; Atick and Redlich [7]), sparse coding (Olshausen and Field [8]) and predictive coding (Rao and Ballard [9]), recovering localized, oriented, bandpass filters that resemble the measured ones. What neither line fixes is how far the population of receptive fields extends along any one degree of freedom, and it is those widths that we measure here.

The receptive fields in the primary visual cortex can be seen as a front-end upon which later stage vision processes strongly depend. Understanding the computational functions of these receptive fields is therefore essential if one aims at understanding the computational functions in the vision system. According to the taxonomy by Hubel and Wiesel [10–13], there are two main types of visual neurons in the primary visual cortex in terms of simple cells and complex cells, where simple cells are often modelled as linear, while the complex cells are genuinely non-linear. In this paper, we focus primarily on simple cells, motivated by their property of being assumed to be simpler to model.

For the purpose of understanding these visual neurons, explicit reconstructions of the receptive fields of simple cells have been performed by DeAngelis et al. [14, 15], Ringach [16, 17], Conway and Livingstone [18], Johnson et al. [19], Walker et al. [20] and De and Horwitz [21]. Functional properties of such neurons have also been characterized in terms of their orientation selectivity properties by Watkins and Berkley [22], Rose and Blakemore [23], Schiller et al. [24], Albright [25], Ringach et al. [26], Nauhaus et al. [27], Scholl et al. [28], Sadeh and Rotter [29], Goris et al. [30], Sasaki *et al*. [31] and Almasi et al. [32], with a conceptual theoretical bridge between these complementary ways of characterizing visual neurons established in Lindeberg [33].

Computationally, the functional properties of simple cells have mainly been modelled according to two different schools of computational models, either in terms of Gabor functions by Marcelja [34], Jones and Palmer [35, 36], Porat and Zeevi [37], Ringach [16, 17], Citti and Sarti et al. [38], Serre et al. [39], Sarti et al. [40], Cocci et al. [41], Barbieri et al. [42], Baspinar et al. [43, 44] and De and Horwitz [21], or in terms of Gaussian derivatives by Koenderink and van Doorn [45–47], Young [48], Young et al. [49, 50] and Lindeberg [51–53]. Theoretical models of early visual processes in terms of Gaussian derivatives have specifically been formulated by Lowe [54], May and Georgeson [55], Hesse and Georgeson [56], Georgeson *et al*. [57], Hansen and Neumann [58], Wallis and Georgeson [59], Wang and Spratling [60], Pei *et al*. [61], Ghodrati *et al*. [62], Kristensen and Sandberg [63], Ruslim *et al*. [64] and Wendt and Faul [65].

Four things happen to the image on the retina as an animal and an object move relative to one another: (i) The object comes nearer or moves further away, and its image grows or shrinks: a spatial scaling transformation. (ii) The surface being viewed slants away, and its image is compressed along one axis while the perpendicular axis is left alone: a spatial affine transformation. (iii) The object moves across the visual field, so that its image traces a slanted path through space and time: a Galilean transformation. (iv) And the same event unfolds quickly or slowly: a temporal scaling transformation (Fig 1A). According to the normative theory of visual receptive fields proposed in Lindeberg [51–53] and substantially extended in Lindeberg [66–69], the simple cells of the primary visual cortex form a basis of receptive fields that is covariant under these four classes of geometric image transformations. Covariant here means that when the image is transformed, the response of one receptive field is reproduced by another receptive field elsewhere in the population, so that the two views can still be brought into correspondence. Covariance in this sense constrains what the population has to contain. To cover the degrees of freedom of the four classes of geometric image transformations, the shapes of the receptive fields have to vary along seven parameters: (i) the spatial size (spatial scale), (ii) the preferred orientation in image space, (iii) the degree of elongation, (iv) a fourth spatial degree of freedom, (v) the preferred direction in joint space-time, (vi) the preferred velocity in joint space-time and (vii) the temporal duration (temporal scale), see Lindeberg [67, 68] for more detailed motivations.

**Fig 1.**
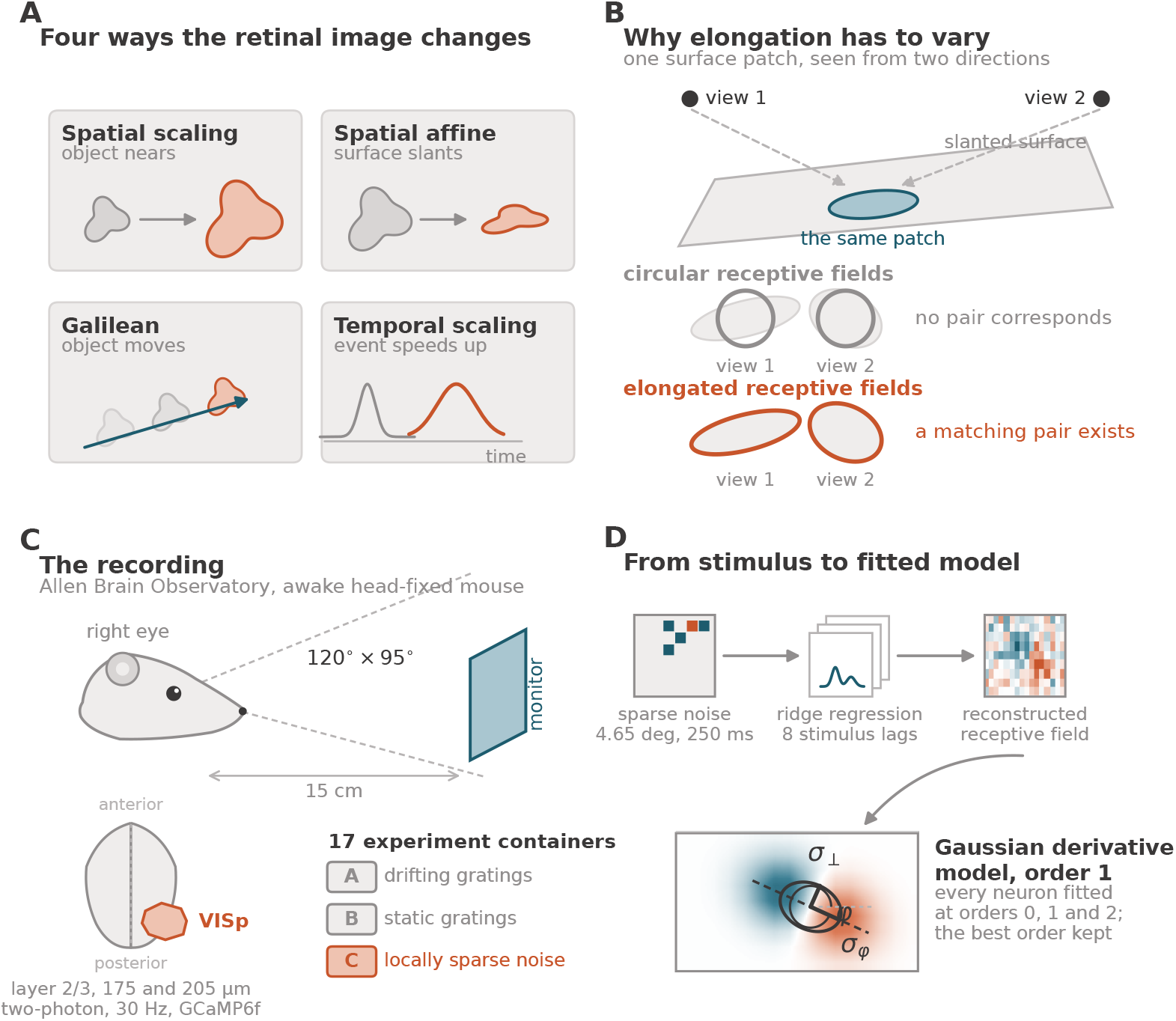
Framework, dataset and pipeline. (**A**) The four geometric image transformations under which the receptive field family is covariant. (**B**) Why the population must vary in elongation: a single surface patch seen from two directions projects to differently shaped image regions, and only a population containing elongated fields supplies a matching pair. (**C**) The Allen Brain Observatory recording: stimuli are presented monocularly to the right eye on a monitor spanning 120^*°*^ *×* 95^*°*^, and layer 2/3 of VISp is imaged across 17 experiment containers with three stimulus sessions each, with receptive fields mapped from the locally sparse noise stimulus of session C. (**D**) The reconstruction pipeline, from a locally sparse noise frame through reverse correlation to a fitted Gaussian derivative model.

These predictions come from a normative analysis, which asks what receptive fields an idealized visual system ought to have if one common front end is to serve many different later processes, rather than fitting a descriptive model to what the recordings happen to show. The requirements it starts from are structural rather than statistical: (i) The earliest layers should be linear. (ii) The same operation should apply everywhere in the visual field. (iii) Representations at neighbouring scales should be consistent with one another, in the sense that smoothing twice by small amounts gives exactly what a single smoothing would give, at the scale whose *variance* is the sum of the two. (iv) And a coarser representation should never invent structure that was absent at finer scales, so that the value at a local maximum cannot rise as the scale increases. From these requirements, the family of affine Gaussian derivatives follows by necessity, with no appeal to the statistics of the images the animal happens to see (Lindeberg [53]).

The same distinction is now familiar in machine vision. A convolutional network shares its weights across the image, which builds covariance under translations into the architecture rather than leaving it to be learned from examples. Group-equivariant networks (Cohen and Welling [70]) extend that construction to rotations and scalings for the same reason: a representation that transforms with the image can be matched across views without the network having to encounter every view during training. Networks built directly from Gaussian derivatives make the connection to the present theory explicit (Lindeberg [71]). The contrast with the statistics-driven accounts is the same in both fields: the covariance is either designed in, or it has to be learned from whatever images the system is exposed to.

What the covariance buys can be seen with a slanted surface. A receptive field, that is circular in the image, samples an elliptical patch of that surface, and the shape of the ellipse changes with the viewing direction. Two views of the same patch are then measured over regions that do not correspond, and any shape cue computed from the two responses carries a systematic error. A population that also contains elongated receptive fields, at a range of elongations and orientations, can supply a pair whose backprojections onto the surface do match, and the error disappears (Fig 1B). This is why the theory needs the receptive fields to vary in elongation, and it is the degree of freedom we measure here.

As described in more detail in Lindeberg [67, 68], there is partial support for these predictions from neurophysiological recordings in the primary visual cortex of higher mammals such as primates and cats. The publicly available reconstructions of receptive fields from such recordings are, however, very sparse.

The subject of this paper is to investigate if we could establish support for a subset of these properties by analysing the results of neurophysiological recordings from visual neurons in mice, for which there are substantially more publicly available data. Concerning mice, it should first of all be noted that the visual system of mice is far less developed than the vision system of higher mammals, see *e*.*g*. Huberman and Niell [72] for an overview, and Abballe and Asari [73] for the statistics of natural images taken from the mouse’s own viewpoint. The spatial receptive fields of mouse simple cells were characterized by Niell and Stryker [74], who reported that the Gabor shape parameters of mouse receptive fields overlap those measured in the macaque by Ringach [16], with the difference that mice lack receptive fields with many subfields; see Niell [75] for a review. Using two-photon calcium imaging, as we do here, Bonin *et al*. [76] found linear, simple receptive fields among excitatory neurons in mouse visual cortex, while inhibitory neurons summated non-linearly and were weakly tuned for orientation. Furthermore, recent results by Fu *et al*. [77] suggest that complex spatial features emerge earlier in the visual pathway of mice compared to primates. Thereby, the receptive fields may be more complex than a Gabor or Gaussian derivative model, as used in the theoretical model for stating the predictions in Lindeberg [67, 68]. Hence, it is not at all a priori clear to what extent the predictions about the seven types of variabilities in the shapes of simple cells would hold for mice.

Of these seven variabilities, four are purely spatial and three arise only in joint space-time. Here we address three of the spatial ones, and establish the existence of (i) a moderate variability over spatial scales, (ii) a uniform variability over the preferred orientation and (iii) an explicit variability over the degree of elongation. The three space-time variabilities call for a spatio-temporal analysis of a stimulus that drives the cells over time, which the receptive field mapping used here does not provide; we return to this in the Discussion.

While a uniform variability in the preferred orientation of neurons in the primary visual cortex of higher mammals could straightforwardly be predicted from existing neurophysiological evidence about orientation maps by Bonhoeffer and Grinvald [78], Blasdel [79] Koch et al. [80] and others, the vision system of mice does not have a pinwheel structure. Additionally, while an ideal vision system may according to the general theoretical results in Lindeberg [81] not need to represent more than the finest scale levels in the sensorium, because of cascade smoothing properties from finer to coarser scale levels, the scale levels extracted from our analysis of simple cells are in a rather narrow scale range over roughly one octave. A more notable result that we will present is that we here establish the existence of an explicit variability in the degree of elongation, as was previously predicted for higher mammals in Lindeberg [82] by combining the results of orientation selectivity measurements by Nauhaus et al. [27] and Goris et al. [30] with the theoretical analysis of the orientation selectivity properties of receptive field models in terms of affine Gaussian derivatives in Lindeberg [33], as well as also predicted for genuine spatio-temporal receptive fields in Lindeberg [83], by fitting parameterized idealized models of direction selectivity curves to neurophysiologically measured direction selectivity curves by Livingstone and Conway [84]. From the neurons that we analyzed, this variability in the degree of elongation clearly exceeded the variability that the measurement itself would produce, spanning about a factor of 5. In this way, we provide direct evidence for one of the explicit predictions that follow from the normative theory of visual receptive fields in Lindeberg [53], as developed into the more general set of theoretical predictions in Lindeberg [67].

## Results

### Most of the simple cells are better described by first-order directional derivatives than by higher orders of spatial differentiation

We reconstructed spatial receptive fields for approximately 3,302 Layer 2/3 excitatory neurons across 17 experiment containers. Each neuron was fit to a generalised Gaussian derivative model of orders *m ∈ {*0, 1, 2*}* (Fig 2). Fitting each neuron with its best-fitting order yielded **1**,**017 neurons** that passed the goodness-of-fit criterion (*R*^2^ *≥* 0.3), which we take as the analysed population throughout. Automated order classification assigned 706 of these to *m*=0 (69%), 125 to *m*=1 (12%) and 186 to *m*=2 (18%).

**Fig 2.**
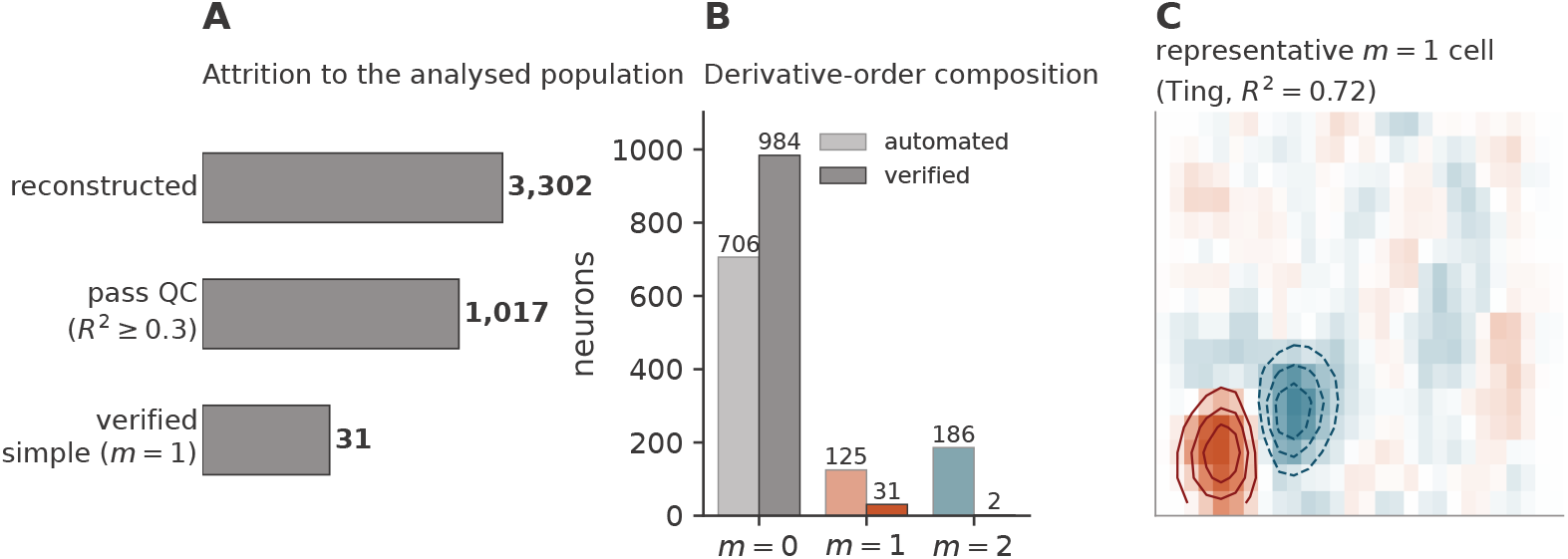
Reconstruction, quality control and derivative-order composition. (**A**) Attrition from approximately 3,302 reconstructed neurons to the 1,017 that pass the goodness-of-fit threshold (*R*^2^ *≥* 0.3, best-order fit) to the 31 verified first-order simple cells. (**B**) Derivative-order composition of the analysed population: the automated assignment (*m*=0: 706; *m*=1: 125; *m*=2: 186) beside the manually verified composition (984 *m*=0; 31 *m*=1; 2 *m*=2). (**C**) A representative verified *m*=1 simple cell, with the fitted model’s iso-response contours overlaid on the smoothed reconstruction.

Automated classification cannot on its own separate genuine first-order simple cells from complex cells or from low-SNR maps overfit to higher orders: an antisymmetric two-lobe profile is a necessary but not a sufficient signature of a first-order simple cell, and complex-cell responses can present with apparent *m ≥* 1 structure under linear reconstruction. We therefore verified every *m ≥* 1 candidate and every borderline case by manual inspection (see Methods), retaining only neurons that could be reasonably confirmed as simple cells. Verification confirmed **31** *m*=1 **simple cells** and only 2 *m*=2 cells, with the remaining maps best described by *m*=0 (Fig 2). Among the neurons we can positively verify as simple cells, first-order directional-derivative structure therefore dominates, outnumbering verified second-order structure by roughly an order of magnitude. This absolute yield is shaped by the spatial resolution of the stimulus (4.65^*°*^/pixel), the temporal integration of the GCaMP6f indicator, the linear receptive-field reconstruction, and a deliberately conservative verification procedure; we consider these constraints in the Discussion.

### The spatial scales vary by about a factor of two

The 31 verified *m*=1 simple cells were distributed across 13 containers (3–5 cells per container, mean 4.2) and both excitatory lines (Slc17a7-IRES2-Cre, *n* = 19; Cux2-CreERT2, *n* = 12). We summarise receptive field size by the geometric-mean scale 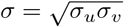, the geometric mean of the scales along and perpendicular to the differentiation axis, which is a single rotation-invariant measure of spatial extent rather than a single-axis width or an arithmetic average. Across the population, the geometric-mean scale spanned 5.37^*°*^ to 12.55^*°*^ (median 7.65^*°*^, mean 7.83^*°*^, standard deviation 1.47^*°*^, coefficient of variation 0.19), a variability of about a factor of two (Fig 3). We take up the interpretation of this scale range in the Discussion.

**Fig 3.**
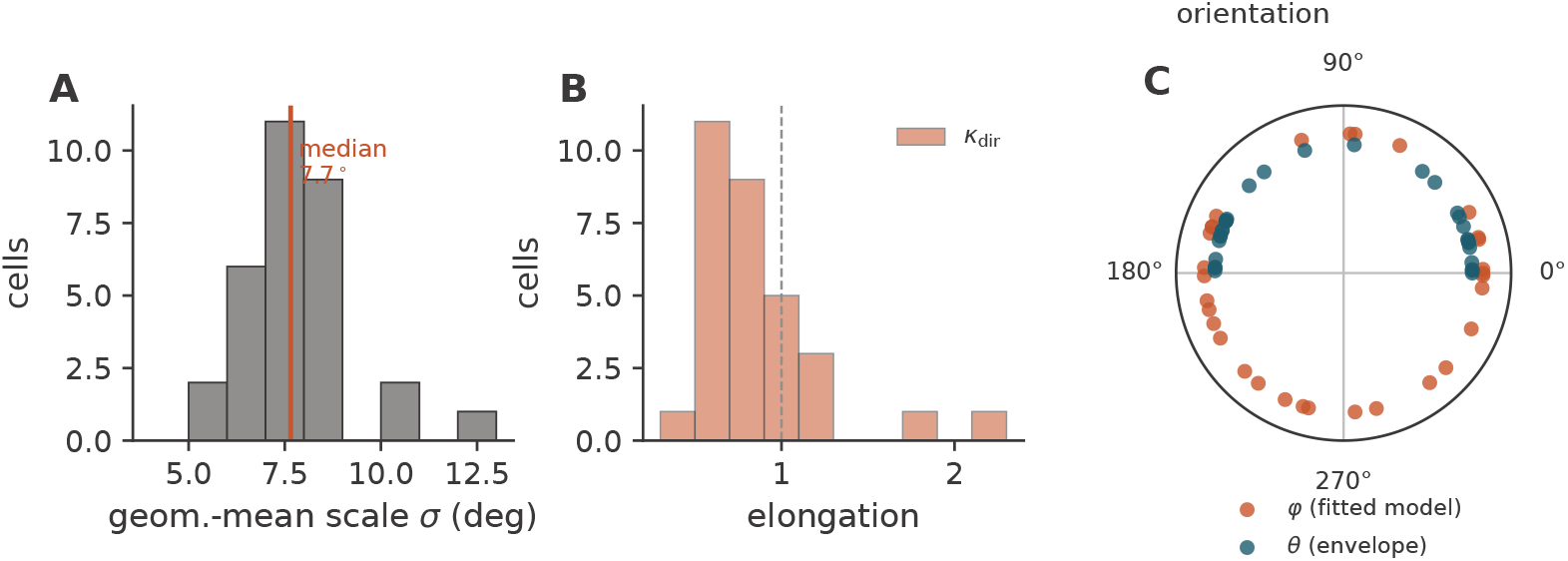
Spatial parameters of the 31 verified simple cells. (**A**) Distribution of the geometric-mean scale *σ* (median marked). (**B**) Distribution of the direction-aware elongation *κ*_dir_ (dashed line at unity). (**C**) Orientation: the differentiation direction *φ* (directional, [0^*°*^, 360^*°*^)) is near-uniform, whereas the long axis of the envelope *θ* (an axis, [0^*°*^, 180^*°*^)) leans towards horizontal.

### The simple cells span a substantial variability in the degree of elongation

Elongation was measured with a direction-aware ratio that takes values both below and above unity, so that its *sign* relative to the differentiation axis is retained: *κ*_dir_ *<* 1 indicates elongation *along* the differentiation axis and *κ*_dir_ *>* 1 elongation *perpendicular* to it. Rather than clustering near a single shape, the simple cells spanned a broad range of elongations: *κ*_dir_ ran from 0.41 to 2.15 across the population (Fig 3; Fig 4). This variability in the degree of elongation is the central positive result of the purely spatial analysis, and it is precisely one of the predictions of the underlying receptive field theory (see Discussion).

**Fig 4.**
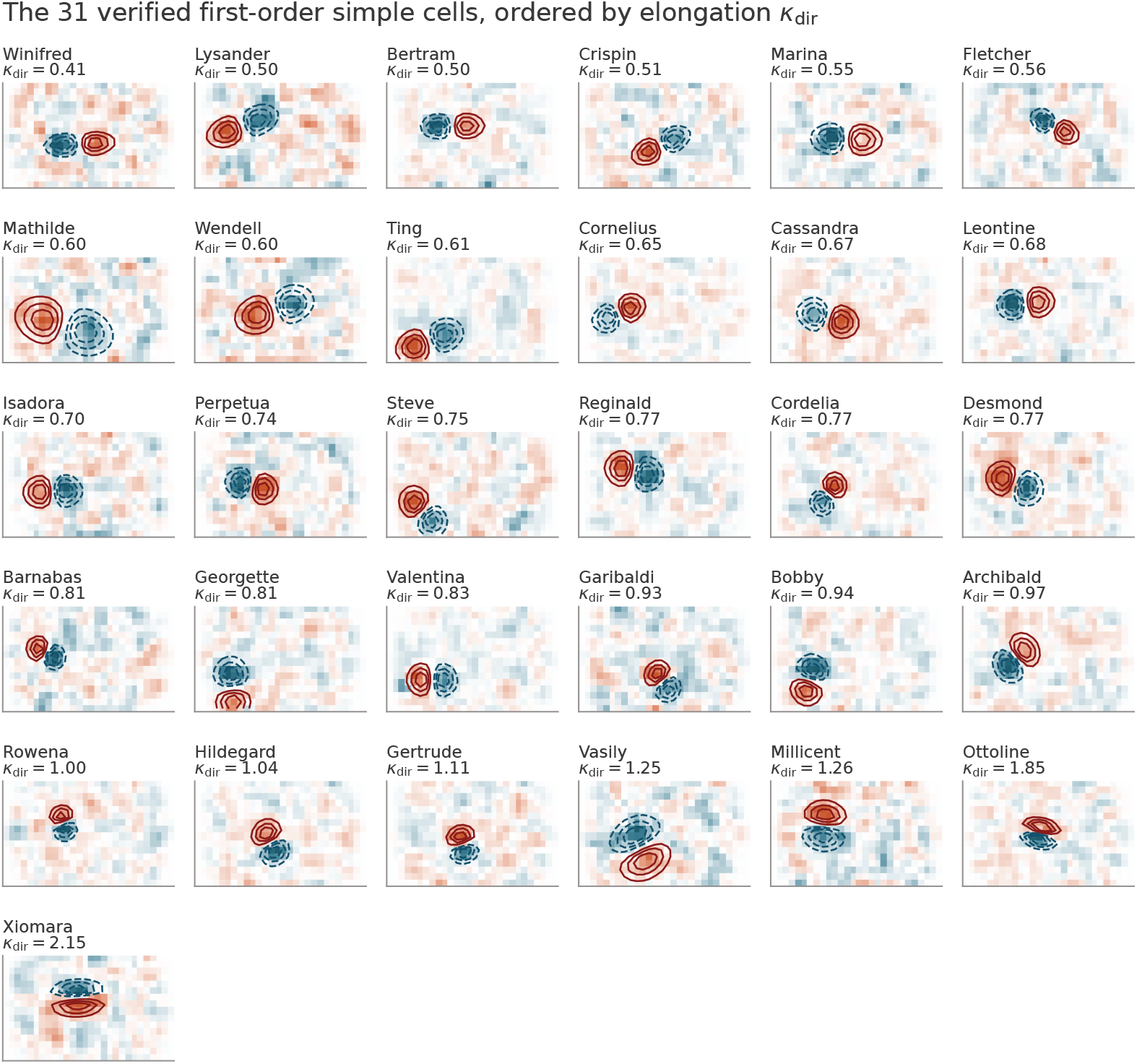
The 31 verified first-order simple cells, ordered by elongation. Each panel shows the smoothed receptive field reconstruction with the fitted *m*=1 model’s iso-response contours overlaid (warm lines, ON lobe; cool lines, OFF lobe), labelled with the cell name and its direction-aware elongation *κ*_dir_. Cells are ordered by *κ*_dir_, so the spread in elongation reads across the array.

This direction-aware measure must be interpreted with care: in the *m*=1 model, *σ*_*φ*_ jointly encodes the size of the oval outline within which the receptive field has any sensitivity at all (the envelope) and the half-separation between the two lobes, so it systematically inflates the apparent extent along the differentiation axis and biases *κ*_dir_ below unity. A separation-free single-lobe control (*κ*_lobe_) reduced the fraction of cells with sub-unity elongation from 24/31 to 14/31, confirming that part — though not all — of the along-axis elongation reflects lobe geometry rather than the underlying envelope. The spread in elongation is not removed by this control: *κ*_lobe_ still ranges from 0.43 to 1.84, a factor of 4.3, against a factor of 5.2 for *κ*_dir_.

The spread in elongation is not an artefact of the measurement, as a control study on the reconstructed maps confirms. For each of the 31 cells we re-embedded a round first-order receptive field (*κ*_dir_ = 1) of the cell’s own size, position, orientation and signal energy in noise taken from the signal-free lags of the cell’s own reconstruction, and refitted it with the unchanged pipeline (see Methods); the reconstruction reproduces the stored maps to within 5 *×* 10^*−*8^ of their peak amplitude, so the control operates on the same maps as the main analysis. Planted elongations were recovered without bias (median ratio of recovered to planted *κ*_dir_ 1.00, interquartile range 0.98 to 1.00). Across the 31 cells the interquartile ratio of *κ*_dir_ was 1.59, whereas round cells measured under the same noise gave 1.24, with 95% of null populations below 1.37 (*p <* 0.001); the median absolute deviation gave the same result (*p <* 0.001), and roughly three quarters of the between-cell variance in elongation is attributable to the cells rather than to the measurement. Drawing each cell’s perturbing noise from the other cells rather than from its own reconstruction changed none of this (interquartile ratio of the null 1.25; per-cell interval widths within 0.5%), ruling out any contribution from reusing a cell’s own noise. The observed spread in *κ*_dir_ is therefore significantly greater than the uncertainty in the individual estimates: the dispersion across cells (standard deviation of log *κ*_dir_ 0.37) is about twice the typical per-cell uncertainty (median half-width 0.18), and the intervals of the least and most elongated cells remain separated by a factor of 3.6. The fits do have a heavy tail, with about 5% of refits of a round field returning an elongation more than twofold from unity, so the full range of 0.41 to 2.15 is reported descriptively and the evidence for variability rests on the dispersion of the population as a whole. The spread is also robust to the inclusion criteria: restricting to cells with *R*^2^ *≥* 0.4 (26 cells) or removing the three cells whose first-order fit exceeded the zero-order fit by less than Δ*R*^2^ = 0.05 (28 cells) left the range unchanged, and removing the seven cells whose first-order assignment is marginal against the second-order model left both the range (5.2) and the interquartile ratio (1.67) intact, since none of these cells is among the extremes. Per-cell estimates with bootstrap intervals are given in S1 File and S3 Table, and the full analysis in S2 Fig.

The smoothing applied before fitting biases the measured elongations in a single direction, which makes the fitted spread a lower bound on the true one. The maps are smoothed by 0.75 pixels (3.49^*°*^) before fitting, so what the fit returns is the scale of the smoothed map. Because Gaussian smoothings compose by adding variances, 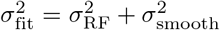 on both axes, and adding the same quantity to the numerator and the denominator pulls any ratio toward unity — most strongly for the smallest receptive fields. Removing that contribution in quadrature gives a corrected 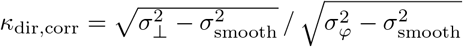, which ranges from 0.29 to 2.67, a factor of 9.1. All 31 cells remain above the smoothing scale on both axes, so the correction is defined throughout. We report both because they answer different questions: *κ*_dir_ is what the measurement delivers, and *κ*_dir,corr_ is the first-order estimate of the underlying receptive field. Either way the conclusion is the same, and the corrected figure makes the direction of the bias explicit: the true spread is wider than the fitted one, not narrower. This argument concerns the smoothing alone; measurement noise acts on the extremes of the distribution in the opposite direction, which is why the variability is assessed against the noise floor above rather than by its raw range.

### Scale and elongation show no detectable coupling among the simple cells

Scale and elongation are predicted to vary independently, as two separate degrees of freedom. Among the 31 verified simple cells we found no significant association between them (*κ*_dir_ against *σ*, Pearson *r* = *−*0.14, *p* = 0.45; symmetric *κ* against *σ, r* = 0.18, *p* = 0.34). A positive correlation is present across the full order-corrected population (*r* = 0.52, *n* = 1,017), but this pools neurons of different types and is in part generated by the isotropic pre-smoothing, which recovers small receptive fields as rounder than they are; we therefore do not interpret it as a property of the simple cells and report it, with the smoothing-floor analysis, only in S4 Text.

### Orientation is uniformly distributed at the population level

The differentiation direction *φ* of the fitted first-order model, defined over [0^*°*^, 360^*°*^) and measured counter-clockwise from rightward horizontal, was not significantly different from uniform across the 31 simple cells (mean resultant length *R* = 0.13, Rayleigh *p* = 0.60; Fig 3), matching the predicted uniform variability over the preferred orientation in image space. The direction was well determined: re-embedding each fitted model in its own measurement noise gave a median 68% interval of 6^*°*^ (largest 12^*°*^). The long axes of the receptive field envelopes, by contrast, leaned towards horizontal (25 of 31 within 30^*°*^). Refitting each cell’s geometry planted at uniformly random orientations in its own noise produced no such lean, so it is introduced neither by the fitting procedure nor by the measurement noise; whether it reflects the stimulus or the cells themselves cannot be resolved from these data.

### Receptive field parameters show no spatial clustering

To test for global topological organisation, we computed the spatial autocorrelogram of elongation across cortical position. On the full order-corrected population — the appropriate basis for this test, since the *m*=1 subset is too sparse to support a standalone autocorrelogram — there was no spatial structure: the correlation between pairwise cortical distance and pairwise elongation difference was effectively zero (*r* = *−*0.004 over 28,510 neuron pairs across 17 containers; Fig 6). Because pairs that share a neuron are not independent, significance was assessed by permuting neuron labels over cortical positions within each container rather than by the parametric *p*-value, which gave *p* = 0.80. Neighbouring neurons were no more alike in elongation than distant ones (pairs closer than 50 µm, one-sided *p* = 0.32), and only one of the 17 containers showed a nominally significant correlation, about as many as expected by chance. Receptive field scale likewise showed no cortical gradient (*p* = 0.89), with a weak tendency for neurons closer than 50 µm to have more similar sizes (one-sided *p* = 0.04) that does not survive correction for the four tests. The 31 simple cells likewise showed no obvious within-or across-container parameter clustering when plotted in cortical space. Receptive field shape in mouse VISp is therefore organised in a salt-and-pepper fashion, without a global spatial map.

### Similar characterisations in the monocular and the binocular zones

We compared receptive field scale and elongation between the frontal/nasal binocular zone and the monocular zone, classified by receptive field centre azimuth (Fig 6; see Methods). Neither measure differed detectably between the zones (binocular *n* = 13, monocular *n* = 18; *σ*: Mann–Whitney *p* = 0.17, rank-biserial correlation *−*0.30, 95% CI *−*0.69 to 0.10; *κ*_dir_: *p* = 0.67, rank-biserial correlation *−*0.09, 95% CI *−*0.50 to 0.33; negative values indicate marginally smaller values in the binocular zone). We discuss below why this similarity does not by itself establish an absence of binocular specialisation.

### Goodness of fit does not separate the Gaussian derivative from the Gabor model

The Gabor parametrisation, introduced by Marcelja [34] and fitted to simple cells by Jones and Palmer [35, 36], remains the most widely used description of simple cell receptive fields, although its adequacy has been questioned before: Hawken and Parker [85] found difference-of-Gaussian models superior to both the Gabor function and the second derivative of a Gaussian for spatial frequency tuning in monkey striate cortex, and Stork and Wilson [86] argued that the theoretical and empirical case for Gabor functions is weaker than usually assumed. We therefore tested on our own maps whether the two families can be told apart by how well they fit.

They cannot. On the 31 verified simple cells, fitting both families from the same 16 starting points, the Gabor reached a median *R*^2^ of 0.50 against 0.48 for the first-order Gaussian derivative, fitting marginally better in 31 of the 31 cells, and the Akaike Information Criterion favoured the Gabor in 30 of them despite its two extra parameters (median ΔAIC = +19.4; Fig 7). The comparison is a fair one. Both models describe the receptive field as an oval region of the visual field, free to be long in any direction (an anisotropic Gaussian envelope), and in both the lobe structure inside that region is laid out along the oval’s own long axis rather than at some angle to it. Neither model, in other words, is given a shape the other cannot make (Methods). The two models are close enough over the range of shapes these cells occupy that the ordering between them turns on how each is optimised rather than on which is right: fitting the Gabor from a single warm start, as is easily done, costs it enough *R*^2^ to reverse the comparison. We therefore make no claim from goodness of fit in either direction, and rest the choice of parametrisation on the structural grounds set out in the Discussion. This does not weaken the shape estimates reported above, which depend on the Gaussian derivative model being an adequate description of these maps, and it is; it removes only the separate claim that it is the better-fitting one.

## Discussion

Using the generalised Gaussian derivative model, we characterised the spatial receptive fields of Layer 2/3 excitatory neurons in mouse VISp. Our conclusions rest not on the absolute number of simple cells — which, as we discuss below, is a lower bound set by the resolution of the recordings — but on the *structure* of the parameter space among the cells we can positively resolve as simple cells. We are deliberately cautious about the large fraction of maps best fit by *m*=0: because linear reconstruction from sparse noise cannot distinguish a genuine zero-order simple cell from a complex cell, or from a higher-order mechanism that the experiment does not resolve; this fraction should not be read as evidence that the underlying computation is low-order. What we can say concerns the verified simple cells, and here the picture is consistent across every measure: the differentiation directions are uniformly distributed, the spatial scales span about one octave, and — most informatively — the degree of elongation varies substantially. Neither receptive field shape nor binocularity shows a global gradient across cortical position, although local structure below the scale of our sampling, such as the ON/OFF domains described by Tring et al. [87], is not excluded.

The central result is the substantial variability in the degree of elongation. This was not the outcome originally anticipated for this project. The working hypothesis had been that the receptive fields of mouse simple cells would span only two degrees of freedom, spatial scale and orientation, on the grounds that the visual ecology of the mouse, with laterally placed eyes, low acuity and limited binocular overlap, might be served by so restricted a repertoire. A variability over the degree of elongation is a third degree of freedom, and the simple cells span it well beyond the noise floor of the measurement, over roughly a factor of five. This extends to the mouse a prediction advanced for higher mammals in Lindeberg [82] by combining the orientation-selectivity measurements of Nauhaus et al. [27] and Goris et al. [30] with the theoretical analysis of orientation selectivity for Gaussian-derivative receptive fields in Lindeberg [33], that the primary visual cortex should span an explicit variability over the degree of elongation. Establishing that this variability is present in the mouse, a species without an orientation-map architecture, shows that a variability over the affine (elongation) degree of freedom does not depend on a pinwheel organisation to be expressed — it is carried in a salt-and-pepper arrangement instead. This is a stronger and different claim than the reduced-repertoire hypothesis we set out with, and it places the affine covariance property of the normative theory (Lindeberg [53, 67]) on a firmer empirical footing.

The comparatively narrow range of spatial scales — about one octave — should be read in this light rather than as a further sign of a reduced repertoire. According to the cascade smoothing property of the scale-space framework (Lindeberg [81]), an ideal visual system need not represent a wide range of scales explicitly in the sensorium: it can represent the finest scale levels and derive coarser-scale responses by successive smoothing. A limited spread of directly represented scales is therefore consistent with, rather than contrary to, the theory. Two caveats reinforce this reading. First, the reconstruction is driven by a stimulus of fixed element size (4.65^*°*^), which sets the band of scales the experiment can excite; receptive fields substantially larger or smaller than this element may be under-represented, so the observed scale range is likely an underestimate of the true range, and stimuli spanning several element sizes would be needed to measure it properly. The direction of that bias can be stated more precisely. Every smoothing stage in the measurement chain, from the finite stimulus element to the pre-smoothing applied before fitting, combines with the receptive field itself so that variances add, and what is recovered has a scale of 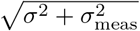 rather than *σ*. A population spanning a wide range of true scales at or below *σ*_meas_ is therefore recovered as a population spanning a much narrower range, and a narrow measured range is the expected outcome whether or not the underlying variability is narrow. Here *σ*_meas_ = 3.74^*°*^, from the 0.75-pixel pre-smoothing (3.49^*°*^) and the 4.65^*°*^ stimulus element (1.34^*°*^) combined in variance, which is 70% of the smallest scale we fit. Subtracting it turns the measured 5.37^*°*^–12.55^*°*^ into 3.86^*°*^–11.98^*°*^, widening the range from 1.22 to 1.64 octaves. That correction is first-order and depends on treating each stage as Gaussian, so we give it as a bound on the direction and size of the bias rather than as a replacement for the fitted range. Second, the uniform distribution of differentiation directions, recovered from the fitted model, is the orientation counterpart of the same picture: a uniform spread over orientation with no spatial gradient, as expected of a species whose cortex lacks the pinwheel maps of primates and carnivores.

Several methodological choices were essential to these conclusions and may be useful more broadly. Estimated receptive field shape is model-dependent and should be reported with the model that produced it, since the Gabor carrier concentrates amplitude along the spatial-frequency axis and so returns a different elongation from the same map. Which model to prefer, though, cannot be settled by goodness of fit: the two families approximate one another over the range of shapes that simple cells occupy, and the difference in explained variance between them is smaller than the difference made by how each is optimised, as our matched-start comparison shows. The case rests instead on the families being justified differently and supporting different operations. The Gabor rests on an optimality argument, minimising joint uncertainty in space, spatial frequency and orientation (Daugman [88]); the Gaussian derivatives follow by necessity from the structural requirements set out above. One qualification applies to both families equally. In their full form (the affine Gaussian derivative and the affine Gabor), each lets the lobes of the receptive field run at an angle to the long axis of its oval outline rather than along it, and the two have been analysed together on that basis (Lindeberg [33]). We fit the aligned version of each, with the lobes along the long axis. The comparison is therefore matched, but between two restricted models, and the angled versions remain to be tested against these data. Gaussians compose across scales, in that two successive smoothings equal a single smoothing whose variance is the sum of the two, whereas Gabors do not, so only the former supports the cascade construction we appeal to when interpreting the scale range. Gaussian derivatives of order one and above integrate to zero by construction, while the even-phase Gabor does not unless corrected (Stork and Wilson [86]). And the affine Gaussian derivative family is closed under affine transformations of the image domain, which is the property the present analysis rests on throughout. Orientation estimation for near-isotropic edge detectors likewise requires care: the envelope orientation *θ* is poorly defined when elongation is low and changes meaning with the sign of the elongation, whereas the differentiation direction of the fitted model is fixed by the receptive field’s antisymmetry and remains well determined in that regime.

And because the differentiation-axis scale of an *m*=1 field conflates envelope size with lobe separation, a direction-aware elongation must be read alongside a separation-free control (*κ*_lobe_) before any claim that the field is elongated at all. These considerations generalise to any structured receptive field model fit to maps that are coarse, and more coarsely sampled in one direction than the other (anisotropic sampling).

Several limitations bound these conclusions, most flowing from a single source: the absolute composition of the population is shaped by the measurement as well as by biology. The 4.65^*°*^/pixel stimulus grid may not resolve the fine two-lobe geometry that distinguishes a first-order edge detector from a zero-order blob, so genuine simple cells can be recovered as *m*=0 or fail the goodness-of-fit threshold; this concern is not hypothetical, since Yeh *et al*. [89] found in macaque that layer 2/3 receptive fields mapped with sparse noise collapse to a single, less elongated subregion, while the same cells mapped with Hartley subspace stimuli show several elongated subregions, and that sparse-noise maps in the upper layers are the poorer predictor of grating-derived preferred orientation, so both our *m*=0 majority and our elongation estimates are likely to be shaped by the stimulus ensemble, in a direction that makes receptive fields appear less elongated and so compresses the measured spread rather than widening it; the slow dynamics of GCaMP6f blur responses and lower the signal-to-noise of the reverse-correlation estimates; linear reconstruction from sparse noise is blind to the nonlinear, complex-cell structure common in mouse V1; and our manual verification of derivative order was deliberately conservative. The *∼*3% simple-cell yield is therefore a lower bound, set by several contributing factors rather than by the rarity of simple cells alone. Complex cells, which are the more numerous type in the primary visual cortex, contribute only weakly to a linear reconstruction and are largely absent from the recovered simple-cell set; this alone may account for a severalfold reduction. Beyond cell type, the measurement is selective in shape and in motion. Receptive fields much smaller or much larger than the 4.65^*°*^ stimulus element are poorly excited, and strongly elongated receptive fields span too few sample positions along their narrow axis for the two-lobe signature to survive, so both the smallest and the most elongated cells are preferentially lost or recovered as *m*=0. The slow dynamics of GCaMP6f lower the signal-to-noise of every reverse-correlation estimate, and our verification of derivative order was deliberately conservative. A further factor follows from the theory itself: if the simple-cell population is also expanded over the degrees of freedom of Galilean transformations (Lindeberg [83]), then many cells would be adapted to non-zero image velocities, with proportionally more at higher velocities. Such velocity-adapted cells are slanted in space-time and have no single stationary spatial profile, so the present purely spatial reconstruction, which retains one temporal lag per cell, cannot capture them. The near-absence of *m*=2 cells is correspondingly a grid and reconstruction artefact in the second-moment estimates rather than a biological absence. The verified simple-cell population is also small (*n* = 31), which limits the power of within-*m*=1 analyses — the absence of a scale–elongation coupling within the simple-cell subset, for instance, cannot by itself exclude a coupling too small to detect at *n* = 31 — and the data are restricted to superficial Layer 2/3 excitatory neurons imaged with a single indicator at a single resolution.

One consequence of these measurement limits bears on the range of elongation itself. Modelling of direction-selectivity data from monkey by Lindeberg [83] is best accounted for by receptive fields reaching degrees of elongation of the order of four to eight, whereas we recover no verified simple cell with *κ*_dir_ much above two. The factors above all act in the same direction on the upper tail: the most elongated receptive fields are precisely those hardest to excite and resolve with a fixed-element sparse-noise stimulus, so the measured ceiling is a lower bound on the true one. We therefore cannot distinguish whether mouse simple cells genuinely lack the strongest elongations or whether these are present but beyond the reach of this measurement; resolving this would require higher-resolution mapping or stimuli better matched to elongated, motion-adapted receptive fields.

The binocular comparison is necessarily indirect, since the public dataset presents all stimuli monocularly, to the right eye, and contains no dichoptic or eye-specific conditions. We found similar receptive field scale and elongation in the frontal/nasal binocular zone and the monocular zone. This similarity should not be over-read: an absence of a detectable difference in these two spatial parameters does not establish an absence of binocular specialisation, which could reside in eye-specific properties — disparity tuning, ocular dominance, interocular matching — that this dataset cannot probe. The comparison constrains scale and elongation across the visual field; it does not license a general claim of no specialisation.

These limitations map directly onto the most informative next steps. The present analysis is purely spatial, and within this dataset it has to be: in a first analysis of the drifting-grating responses, the temporal-frequency preferences of nearly all matched cells sat at the lowest frequency presented, and the calcium signal does not preserve the response modulation needed to recover direction and speed. The scale-space framework is fundamentally spatio-temporal: extending the Gaussian derivative model into the temporal domain — using the drifting-grating responses available in the same recordings to estimate temporal scale and velocity tuning — would test whether the variabilities we observe in space are joined by the predicted variabilities over the preferred direction in joint space–time, the preferred velocity, and the temporal scale. Higher-resolution sparse-noise mapping, or deconvolution-based response estimation, would raise the simple-cell yield and allow the *m*=2 question to be revisited free of the grid artefact. A quantitative test of how the elongation variability compares across species requires receptive-field-shape distributions measured under matched conditions in map-bearing species such as primates and carnivores. And dedicated monocular or dichoptic stimulation would turn the indirect binocular comparison into a direct assay of binocular interaction and disparity tuning.

Taken together, our results support a view of mouse primary visual cortex in which the receptive fields of simple cells span the variabilities predicted by the covariance properties of the normative scale-space theory — notably an explicit variability over the degree of elongation — expressed in a salt-and-pepper arrangement rather than through the global orientation maps of foveate, binocular species.

## Materials and methods

### The Allen Brain Observatory Visual Coding dataset

Our analysis is based on the two-photon calcium imaging arm of the Allen Brain Observatory Visual Coding dataset (de Vries *et al*. [90]; Allen Institute for Brain Science [91]), a standardised survey of visually evoked activity in the cortex of the awake mouse. The full survey comprises recordings from approximately 60,000 neurons in six visual areas, at four cortical depths, across twelve transgenic lines expressing the genetically encoded calcium indicator GCaMP6f, obtained from 243 adult mice. Because every recording follows the same protocol, the same instrument calibration and the same stimulus set, receptive fields measured in different animals can be pooled without the between-laboratory differences in stimulus and analysis that complicate comparisons across the older single-unit literature.

During a recording the mouse is awake and head-fixed and performs no task, with body posture and the position of the stimulated eye monitored throughout. Visual stimuli are presented monocularly on a calibrated liquid crystal monitor placed 15 cm from the right eye and spanning approximately 120^*°*^ *×* 95^*°*^ of visual space, gamma corrected to a mean luminance of 50 cd m^*−*2^. Because the monitor is flat and close to the eye, all stimuli are spherically warped before display so that apparent size, speed and spatial frequency are constant across the screen as seen from the mouse. That warping is what allows a stimulus element to be treated as subtending a fixed visual angle wherever it falls on the display, and our use of a single degrees-per-pixel conversion throughout depends on it. Neuronal fluorescence is imaged at approximately 30 Hz, and the released data comprise segmented regions of interest, their extracted fluorescence traces, the corresponding Δ*F/F* traces, a set of deconvolved event times, and frame-by-frame stimulus tables.

The organising unit of the dataset is the *experiment container*, which is one mouse imaged at one cortical depth in one visual area (Allen Institute for Brain Science [92]). Each container holds three imaging sessions of roughly one hour, labelled A, B and C, recorded over the same field of view on separate days, so that a segmented neuron carries a single persistent identifier across all three and can be followed from one stimulus protocol to another. The sessions carry different stimuli: session A presents drifting gratings and two natural movies, session B presents static gratings, natural scenes and a natural movie, and session C presents the locally sparse noise stimulus used here together with two natural movies. For containers acquired after September 2016, the third session was replaced by a variant, labelled C2, in which the locally sparse noise stimulus is split into two blocks at two different element sizes; this distinction determines the spatial resolution at which a receptive field can be mapped, and we return to it below.

### Selection of experiments

We restricted the analysis to excitatory neurons of cortical Layer 2/3 in the primary visual area, VISp, selecting experiment containers from the two Cre driver lines that label this population, Slc17a7-IRES2-Cre, which is pan-excitatory, and Cux2-CreERT2, which is specific to Layers 2/3 and 4, at reported imaging depths between 150 µm and 250 µm (realised depths 175 µm and 205 µm). We further required that all three imaging sessions were present, so that the same neurons could in principle be followed between the receptive field mapping of session C and the drifting gratings of session A, and that receptive field mapping had been carried out at the finer of the two element sizes. These criteria yield **17 complete experiment containers** from which **1**,**017 neurons** passed the goodness-of-fit criterion described below (*R*^2^ *≥* 0.3) and constitute the analysed population.

The data were accessed through the AllenSDK using the BrainObservatoryCache interface. In the terms used by that interface, our recordings are drawn from the targeted structure VISp, the Cre lines Slc17a7-IRES2-Cre and Cux2-CreERT2, the session types three session A and three session C2, and the stimulus locally sparse noise 4deg.

### Receptive field mapping stimulus

Spatial receptive fields were mapped from the Session C locally sparse noise stimulus, in which several white and black squares appear in each frame, each 4.65^*°*^ on a side, no two within 23^*°*^ of one another, with the remainder of the display at mean luminance. We used the original `locally_sparse_noise_4deg` stimulus, whose native template is a 16 *×* 28 grid of 4.65^*°*^ pixels. The reconstruction reads grid dimensions and pixel size directly from the stimulus template, so the pipeline adapts automatically to stimulus resolution; all analysed neurons were recorded under the 4.65^*°*^ variant. The coarser 9.3^*°*^ (locally sparse noise 8deg) stimulus released in 2017 was not used in the final analysis population. Receptive fields were analysed on the native 16 *×* 28 grid at native resolution (one pixel = 4.65^*°*^ of visual angle).

### Receptive field reconstruction

Receptive fields were reconstructed by ridge regression (Hoerl and Kennard [93]) of the signed stimulus against the calcium response. We used the released Δ*F/F* traces throughout and did not use the deconvolved event times, which rest on further assumptions about the indicator and would change what the reconstruction estimates.

Two clocks run through the procedure and are kept distinct below. Stimulus frames are presented for 250 ms each with no intervening blank, and each display location is occupied by a spot on approximately 115 occasions over the session, with the arrangement of the remaining spots differing on every occasion (Allen Institute for

Brain Science [94]). Fluorescence is sampled at approximately 30 Hz, so one imaging frame is approximately 33 ms. Response windows below are specified in imaging frames and stimulus lags in presentations.

For each cell a scalar response per stimulus presentation was obtained by averaging the Δ*F/F* trace over a window of **5 imaging frames** (*≈* 165 ms) beginning **4 imaging frames** (*≈* 133 ms) after stimulus onset. This delay accommodates the combined visual-pathway latency (*∼*25 ms to 50 ms) and GCaMP6f rise time (*∼*45 ms), and the window spans the calcium transient peak (Chen *et al*. [95]).

Each stimulus frame was signed-encoded (ON pixel = +1, OFF pixel = *−*1, grey background = 0) and lagged over **8 successive stimulus presentations** to form a design matrix, so that the design spans approximately 1.75 s of preceding stimulus. That is considerably longer than the GCaMP6f decay, and deliberately so: rather than fixing the response latency in advance, the reconstruction estimates a map at every lag and retains the one carrying the largest peak absolute weight. The response was regressed on this design matrix by ridge regression with the *L*_2_ penalty *λ* selected by cross-validation over a 25-point logarithmic grid spanning 10^*−*2^ to 10^5^. The fitted coefficients were reshaped into a spatio-temporal receptive field (lags *×* height *×* width), and the retained spatial map was taken as the signed receptive field and split into ON (+) and OFF (*−*) subfields for fitting.

### Gaussian derivative model

Following the generalised Gaussian derivative model of visual receptive fields (Lindeberg [53, 67]), each receptive field was modelled as a two-dimensional Gaussian derivative of integer order *m ∈ {*0, 1, 2*}*. In a frame (*u, v*) aligned to the receptive field, with *u* along the differentiation axis and *v* perpendicular according to the rotation transformation

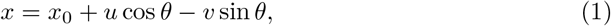

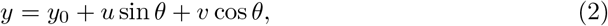

and the Gaussian envelope

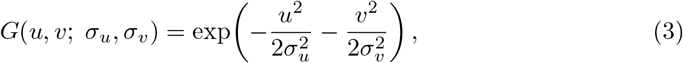

the Gaussian derivative models of orders 0, 1 and 2, respectively, are given by

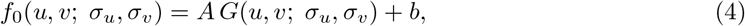

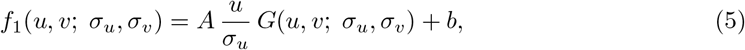

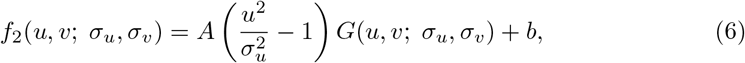

corresponding to a single-lobe blob (*m*=0), an antisymmetric two-lobe edge detector (*m*=1), and a symmetric centre-flank operator (*m*=2); the prefactors are the probabilist’s Hermite polynomials He_*m*_(*u/σ*_*u*_). All the three model share the same set seven free parameters (*A, x*_0_, *y*_0_, *σ*_*u*_, *σ*_*v*_, *θ, b*), where *A* is the amplitude, (*x*_0_, *y*_0_) the center, (*u, v*) the local coordinates in the oriented frame, *θ* the orientation and *b* the offset. Thereby, the order *m* can be compared directly by the goodness of fit. The *m*=1 and *m*=2 models were fit to the full signed map (ON *−* OFF), not to a single dominant subfield, because the zero-crossing between the lobes is the discriminating signature of the derivative order.

The spatial scale is summarised by the geometric-mean scale 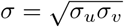 (in degrees of visual angle) and the elongation by *κ*_dir_ = *σ*_*v*_*/σ*_*u*_. The orientation parameter of the model is defined over [0^*°*^, 360^*°*^). For the odd-order model (*m*=1), the sign of the response distinguishes an angle from that angle plus 180^*°*^ (an ON–OFF edge from an OFF–ON edge), so the orientation is genuinely directional and is taken from the fitted model as the differentiation direction *φ* (see *Orientation estimation*). For the even-order models (*m*=0, *m*=2) the response is symmetric and the orientation is an undirected axis, defined over [0^*°*^, 180^*°*^).

### Model fitting

Each order was fit to the signed map by non-linear least squares using the trust-region-reflective solver (Branch *et al*. [96]), which natively handles the box constraints on the parameters. Maps were lightly pre-smoothed with a Gaussian of 0.75 pixel before fitting. To guard against convergence to local minima, we used multi-start optimisation: one moment-based initialisation (from the image moments of |RF| ) plus **15 random initialisations**, retaining the fit with the highest *R*^2^. For *m*=0 (where the Gaussian envelope is non-negative) OFF-dominant maps were sign-flipped before fitting so the amplitude constraint *A ≥* 0 could not collapse the fit; scale and elongation are polarity-invariant; the true polarity was recorded separately and, for *m*=1, sets the sign of the directional orientation *φ*. For each fit we recorded the *R*^2^ improvement of the best start over the moment-based start as a local-minimum diagnostic.

### Noise floor and bootstrap intervals

The ridge regression estimates a spatial map at each of the 8 stimulus lags. Lags beginning at least 1 s after a cell’s best lag carry no stimulus-locked signal but contain the estimator’s noise, with the same stimulus design, regularisation and shrinkage as the best-lag map; lags whose smoothed map correlated with the best-lag map (|*r*| *>* 0.3) were excluded. The retained maps, normalised and augmented by reflection, formed a pool of noise maps, scaled to each cell’s own noise level. For each cell we planted first-order models in this noise and refitted them with the unchanged procedure: a round model (*κ*_dir_ = 1) at the cell’s own geometric-mean scale, the cell’s fitted geometry, and the fitted geometry at uniformly random orientations (100, 100 and 50 draws respectively). Models were planted before smoothing, with each axis reduced in quadrature by the smoothing scale, and scaled so that their energy after smoothing matched the fitted model. The round draws define the noise floor for *κ*_dir_, the draws at the fitted geometry give per-cell bootstrap intervals, and the random-orientation draws test the fitting procedure for an orientation bias. As a control against shared structure between a cell’s template and its own noise, the procedure was repeated drawing each cell’s perturbing noise only from the other cells’ signal-free lags. Population dispersion was compared against 5,000 synthetic populations of 31 round cells, each drawn from its own cell’s noise floor.

### Derivative-order classification and model selection

For each neuron, we fit all three orders and selected the highest-*R*^2^ order subject to a parsimony rule: a higher-order model (*m*=1 or *m*=2) was accepted over *m*=0 only if it improved *R*^2^ by at least Δ*R*^2^ = 0.05; otherwise the simpler *m*=0 model was assigned. This prevents low-SNR maps from being spuriously upgraded by fitting noise. We additionally recorded, per neuron, Δ*R*^2^ relative to the *m*=0 fit. A goodness-of-fit threshold of *R*^2^ *≥* 0.3 was required for inclusion.

### Orientation estimation

The orientation of a first-order receptive field is the direction of its differentiation axis, which is a parameter of the fitted model and remains defined even when the envelope is nearly circular. We took it from the fitted model and resolved its sign from the polarity of the fit, so that it points from the OFF lobe to the ON lobe and is defined over [0^*°*^, 360^*°*^). Angles are reported counter-clockwise from rightward horizontal, as the maps are displayed. The envelope orientation *θ*, the major axis of the fitted Gaussian envelope, is an undirected axis defined over [0^*°*^, 180^*°*^); it coincides with the differentiation axis when *κ*_dir_ *<* 1 and is perpendicular to it when *κ*_dir_ *>* 1. The envelope estimator is biased toward horizontal by the rectangular sampling grid and is used only for the even-order (*m*=0) model; for *m*=1 the differentiation direction is used throughout.

### Direction-aware elongation

The symmetric ratio *κ* = *σ*_major_*/σ*_minor_ *≥* 1 discards the sign of elongation relative to the differentiation axis. To retain it, we instead computed the direction-aware elongation measure *κ*_dir_ = *σ*_*⊥*_*/σ*_*φ*_, where *σ*_*φ*_ is the scale along the differentiation axis and *σ*_*⊥*_ the scale perpendicular to it: *κ*_dir_ *>* 1 indicates elongation perpendicular to the differentiation direction and *κ*_dir_ *<* 1 elongation along it.

### Comparison with a Gabor model

To test whether the goodness of fit distinguishes the two parametrisations, we compared the first-order Gaussian derivative against a nine-parameter 2D Gabor (the seven Gaussian parameters plus the carrier spatial frequency and phase) fit to the same maps. The two are matched in the shapes they can take. In both, the outline of the receptive field is an oval whose length, width and orientation are all free, given by (*σ*_*u*_, *σ*_*v*_, *θ*); this is the most general oval available, since those three numbers specify any covariance matrix (an anisotropic, or affine, Gaussian envelope). In both, the structure inside that oval, the alternating lobes of the Gabor carrier and the positive and negative lobes of the derivative, is laid out along the oval’s long axis *u*. That last constraint is a restriction on both families. Their full forms (the affine Gaussian derivative and the affine Gabor) let the lobes run at an angle to the long axis of the oval, which is the fourth degree of freedom discussed above; we fit the aligned case of each, so the comparison below is between two equally restricted models rather than between the families in full. Because a Gabor started at a high carrier frequency converges to a poor local optimum, and because the Gaussian derivative fits use sixteen starting points, we fit both families from the same sixteen starts, one warm start from the stored parameters and fifteen random. The models were compared by *R*^2^, RMSE and the Akaike Information Criterion, AIC = *n* ln(SS_res_*/n*) + 2*k*, with *k* = 7 and *k* = 9 respectively. This comparison was run on the 31 verified simple cells, for which the reconstructed maps were retained.

### Binocular zone classification

Because stimuli are presented monocularly to the right eye and the dataset contains no dichoptic or eye-specific conditions, eye-specific correlations cannot be measured directly. As an indirect test of binocular specialisation, each *m*=1 simple cell was classified by the azimuth of its reconstructed receptive field centre: cells with centre azimuth *x*_0_ *<* 40^*°*^ were assigned to the frontal/nasal binocular zone and the remainder to the monocular zone. The spatial scale *σ* and elongation *κ* were compared between zones with two-sided Mann–Whitney *U* tests, with the rank-biserial correlation as the effect size and bootstrap 95% confidence intervals.

### Manual verification of derivative order

Because automated order assignment can mislabel low-SNR maps, derivative-order assignments were verified by neuron-by-neuron visual inspection through a purpose-built interactive review interface. For each candidate neuron, the interface displayed the raw reconstructed receptive field and the smoothed map, and, overlaid on each, the fitted parameter contour together with the idealised Gaussian-derivative model rendered at every order (*m*=0, *m*=1, *m*=2), alongside the per-order *R*^2^ statistics. We reviewed every neuron automatically assigned *m ≥* 1, every neuron whose order was ambiguous (a higher-order model exceeding the *m*=0 fit by only a small Δ*R*^2^), neurons whose best fit was weak (*R*^2^ *<* 0.5), and other borderline cases. The complete set of fitted models inspected during this review, with the reconstructed and smoothed maps, the fitted model and its goodness of fit at every order, is provided for all 31 verified cells as S1 File.

Each candidate was judged against criteria we adopted for a visually acceptable Gaussian-derivative fit, following the qualitative comparisons between predicted and measured receptive field profiles in Lindeberg [53]: the number of polarity reversals (lobes) in the model had to match the biological map — one antisymmetric zero-crossing for *m*=1, a symmetric centre–flank pair of zero-crossings for *m*=2 — the fitted envelope had to enclose the receptive field extent without being systematically over- or under-sized, and the differentiation axis had to align with the dominant axis of the map.

Two principles governed the review. Manually confirmed *m*=1 assignments were protected from demotion by any later automated pass. Increased Gaussian smoothing (*σ* = 1.0 and *σ* = 1.5), used as an initialisation aid for the optimiser, was permitted only to recover genuine edge detectors blurred by the coarse grid — that is, to *add* neurons to the *m*=1 set — never to remove already-confirmed ones. This procedure yielded **31 verified** *m*=1 **simple cells**; *m*=2 cells were negligibly rare and excluded from downstream analyses.

Candidate simple cells were reviewed by the first author; the first-order assignment of all 31 verified cells was subsequently confirmed by all three authors from the fitted zero-, first- and second-order models shown in S1 File. Seven cells whose first-order assignment is marginal against the second-order model were identified in this review; removing them changes neither the range nor the dispersion of elongation (see Results), so all 31 are retained.

### Statistical analysis

All tests were two-sided. Dispersion of spatial scale was summarised by the coefficient of variation (CV). The differentiation direction *φ*, defined over [0^*°*^, 360^*°*^), was summarised with standard circular statistics (mean resultant length and the Rayleigh test for non-uniformity). Associations between spatial parameters (e.g. *σ* vs. *κ*) were assessed with Pearson and Spearman correlation, with 95% confidence intervals from the Fisher transformation. Associations between pairwise cortical distance and pairwise parameter difference were tested with Mantel permutation tests (10,000 permutations), permuting neuron labels over cortical positions within each experiment container, because pairs sharing a neuron are not independent.

## Data availability

There are no primary data in this paper. The imaging data analysed in this study are openly available through the Allen Brain Observatory via the AllenSDK (https://portal.brain-map.org/explore/circuits/visual-coding-2p); the specific experiment containers, Cre lines, session types and stimulus are identified in the Methods, so that the exact subset analysed here can be reconstructed without ambiguity. All reconstruction, model-fitting and statistical procedures are described in full in the Methods, permitting independent reimplementation. The fitted receptive field models for the 31 verified simple cells are provided as S1 File. The complete analysis code, together with the derived data sufficient to reproduce every figure and statistic without the raw imaging data, is publicly available at https://github.com/dmescherina/v1-rf-variability-mice under the Creative Commons Attribution 4.0 International (CC BY 4.0) licence, with a versioned snapshot archived at Zenodo (doi:10.5281/zenodo.22883097).

### Software

Receptive field reconstruction and fitting were implemented in Python 3.11 using NumPy, SciPy (`curve_fit`; trust-region-reflective solver), scikit-learn (`RidgeCV`) and pandas, with figures produced in Matplotlib and seaborn; data access used the AllenSDK (`BrainObservatoryCache`). All analyses were run within the project environment.

### Ethics statement

No new animal experiments were performed for this study. All data analysed here were obtained from the Allen Brain Observatory Visual Coding dataset, which is publicly released by the Allen Institute for Brain Science. As reported by the dataset authors [90], all animal procedures for that dataset were approved by the Institutional Animal Care and Use Committee of the Allen Institute for Brain Science. The present work is a secondary analysis of the released imaging data and involved no further procedures on animals.

### Use of AI tools

Claude Code (Anthropic) was used during development of the analysis software to assist with debugging, to refactor working code without changing its results, and to help verify that the pipeline ran end to end in the configuration used for the final computations; this assistance is reflected in the commit history of the public code repository. The AI tool was not used to generate primary data, to perform the statistical analyses, to draw scientific conclusions, or to write the manuscript. All code was reviewed and tested by the first two authors, who retain full responsibility for its correctness; pipeline outputs were validated against fixed reference results, including checks that the reconstructed receptive field maps reproduced the stored canonical maps to within 5 *×* 10^*−*8^ of their peak amplitude and that each fitted model reproduced its recorded goodness of fit. The authors are responsible for all content of this article.

## Supporting information

**S1 File. Fitted receptive field models for the 31 verified first-order simple cells**. For each cell, the raw and smoothed receptive field maps, the fitted first-order model with its differentiation direction, the idealised models of order 0, 1 and 2 with their goodness of fit, and bootstrap intervals for *κ*_dir_ and the differentiation direction.

**S2 Fig. Noise floor for elongation**. Observed *κ*_dir_ for the 31 cells against the range recovered from round receptive fields in each cell’s own measurement noise, the population dispersion against its null distribution, and the recovery of planted elongations.

**S3 Table. Per-cell estimates of scale, elongation and differentiation direction**, with 68% and 95% bootstrap intervals and the per-cell test against a round receptive field.

**S4 Text. Scale–elongation coupling across the full order-corrected population**, with the smoothing-floor analysis and the former Fig 5A.

**Fig 5.**
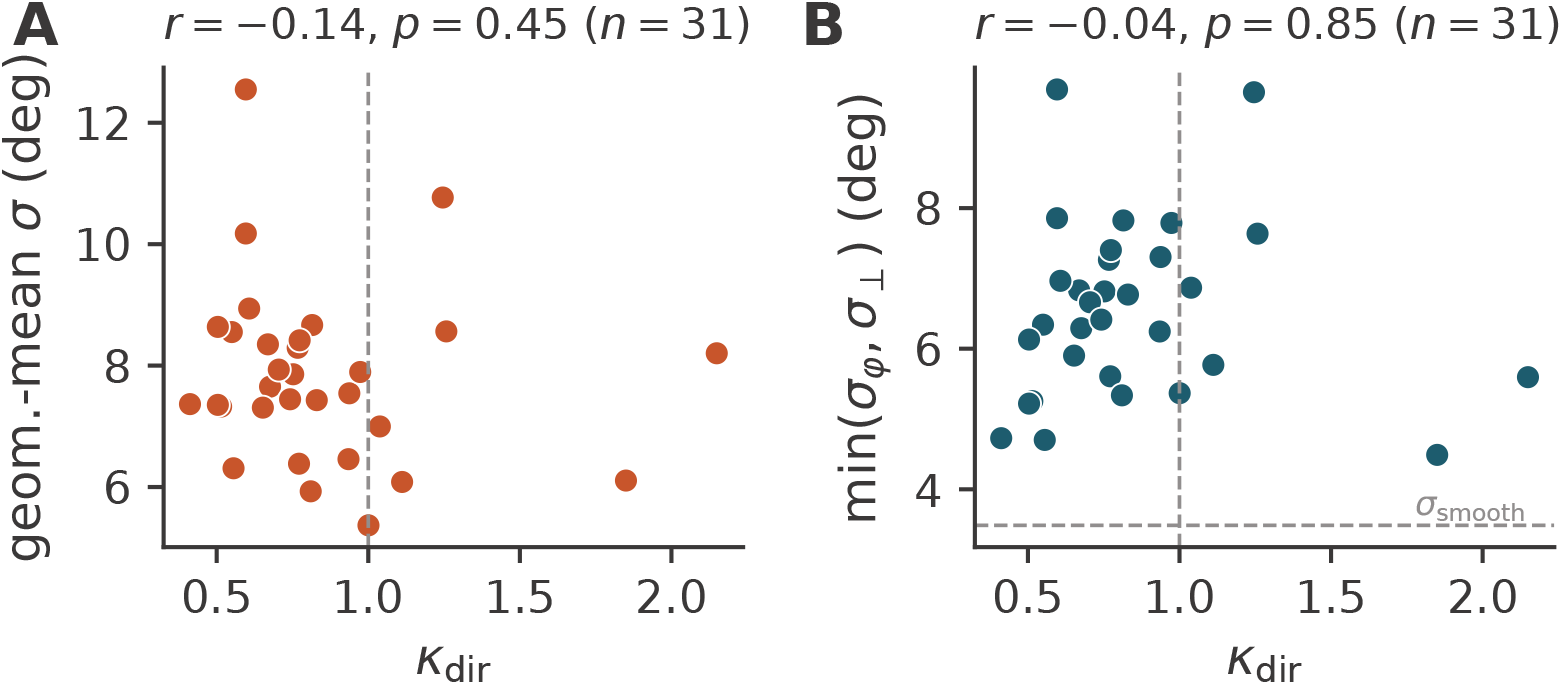
Scale and elongation. Scale *σ* against the symmetric elongation ratio *κ* (**A**) For the 31 simple cells, *κ*_dir_ against the geometric-mean scale *σ*. (**B**) *κ*_dir_ against the minor axis min(*σ*_*φ*_, *σ*_*⊥*_); the dashed line marks the smoothing scale *σ*_smooth_, the floor against which the coupling mechanism can be read.

**Fig 6.**
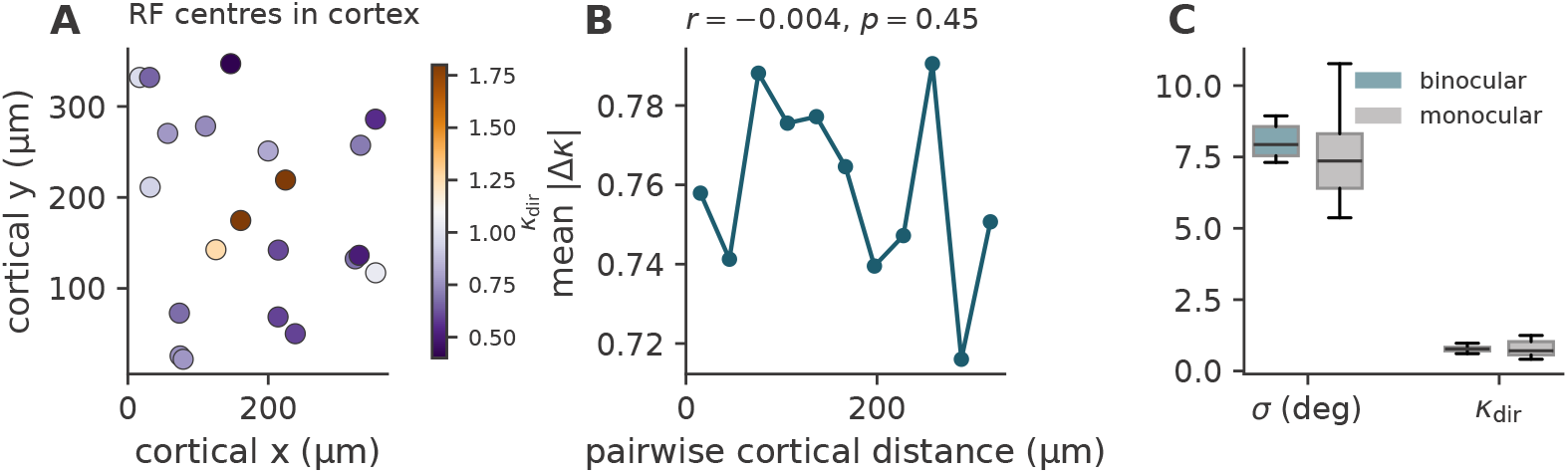
Spatial organisation. (**A**) Receptive field centres of the 31 simple cells in cortical space, coloured by elongation *κ*_dir_. (**B**) Spatial autocorrelogram of elongation on the order-corrected population: mean pairwise elongation difference against pairwise cortical distance, showing no spatial structure. (**C**) Receptive field scale and elongation in the binocular and monocular zones; neither differs detectably.

**Fig 7.**
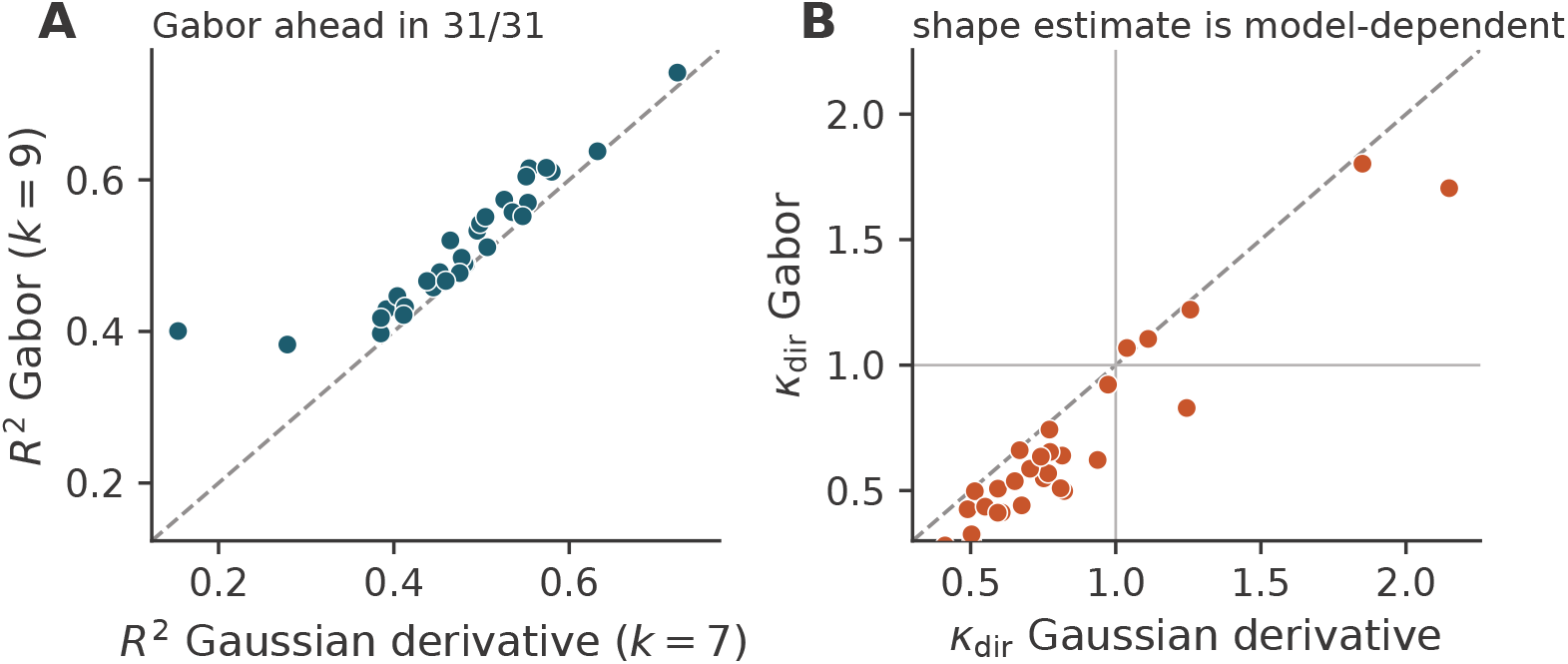
Gaussian derivative versus Gabor. (**A**) Explained variance for the two families under matched optimisation, fitted to the same 31 maps from the same starting points; the two lie close to the identity line. (**B**) Direction-aware elongation *κ*_dir_ recovered by each family from the same maps: the Gabor returns systematically lower values, but the ordering of the cells is preserved (Spearman *ρ* = 0.88, *n* = 30, excluding one cell where the Gabor fit did not converge; *ρ* = 0.79 for all *n* = 31).

