## Supplementary material for "Variabilities in the shapes of the receptive fields of simple cells in mice": S1 File

### Supplementary material for "Variabilities in the shapes of the receptive fields of simple cells in mice"

Fitted receptive field models for the 31 verified first-order simple cells

#### Abstract

This supplementary file accompanies the article Meshcherina, Auffarth and Lindeberg (2026) *Variabilities in the shapes of the receptive fields of simple cells in mice*. It documents the manual verification of derivative order underlying the 31 first-order ($m$=1) simple cells analysed in the main body of the paper. For each verified cell, it presents the reconstructed and smoothed receptive-field maps together with the fitted generalised Gaussian-derivative models for every order ($m$=0, $m$=1 and $m$=2) and their associated goodness of fit, exactly as inspected during the neuron-by-neuron review described in the Methods section. The gallery therefore provides the visual evidence supporting the classification of these cells as first-order simple cells, and allows the fitted receptive-field shapes reported in the main body of the paper to be assessed directly against the underlying reconstructions.

Code and derived data: github.com/dmescherina/v1-rf-variability-mice
Archived at Zenodo: doi:10.5281/zenodo.22883097

### S1 File. Fitted receptive field models for the 31 verified first-order simple cells

Each row shows one cell. From left: (1) the raw ridge-regression map; (2) the same map smoothed at 0.75 pixel, as fitted, with the fitted first-order model's iso-response contours (solid: ON lobe; dashed: OFF lobe) and an arrow along the differentiation direction, pointing from the OFF to the ON lobe; (3-5) the idealised Gaussian derivative models of order 0, 1 and 2 fitted to the same map, with their $R^2$.

The first-order model is the stored canonical fit; the zero- and second-order models are refitted with the same procedure and labelled with their recorded goodness of fit.

Angles are in degrees, counter-clockwise from rightward horizontal as displayed. $\varphi$ is the differentiation direction of the fitted model on [0°, 360°). $\sigma$ is the geometric-mean scale, and $\kappa\_dir = \sigma_\perp/\sigma_\varphi$ (values below 1: elongated along the differentiation axis).

Intervals come from a parametric bootstrap: the cell's fitted model is re-embedded in noise taken from signal-free lags of its own reconstruction and refitted with the same procedure (100 draws). The bracket after κ_dir is the central 68% interval (16th to 84th percentile), and the value after φ is the 68th percentile of the orientation error. 68% intervals are shown because about 5% of refits converge to an alternative solution, which dominates the tails; 95% intervals are given in the accompanying table.

Cells are ordered by κ_dir, as in Fig 4.

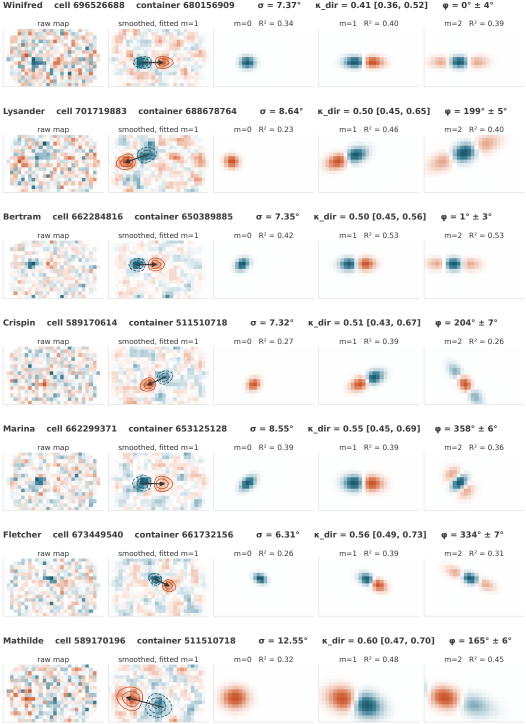
Winifred cell 696526688 container 680156909 σ = 7.37° κ_dir = 0.41 [0.36, 0.52] φ = 0° ± 4°
raw map
smoothed, fitted m=1
m=0 R² = 0.34
m=1 R² = 0.40
m=2 R² = 0.39
Lysander cell 701719883 container 688678764 σ = 8.64° κ_dir = 0.50 [0.45, 0.65] φ = 199° ± 5°
raw map
smoothed, fitted m=1
m=0 R² = 0.23
m=1 R² = 0.46
m=2 R² = 0.40
Bertram cell 662284816 container 650389885 σ = 7.35° κ_dir = 0.50 [0.45, 0.56] φ = 1° ± 3°
raw map
smoothed, fitted m=1
m=0 R² = 0.42
m=1 R² = 0.53
m=2 R² = 0.53
Crispin cell 589170614 container 511510718 σ = 7.32° κ_dir = 0.51 [0.43, 0.67] φ = 204° ± 7°
raw map
smoothed, fitted m=1
m=0 R² = 0.27
m=1 R² = 0.39
m=2 R² = 0.26
Marina cell 662299371 container 653125128 σ = 8.55° κ_dir = 0.55 [0.45, 0.69] φ = 358° ± 6°
raw map
smoothed, fitted m=1
m=0 R² = 0.39
m=1 R² = 0.39
m=2 R² = 0.36
Fletcher cell 673449540 container 661732156 σ = 6.31° κ_dir = 0.56 [0.49, 0.73] φ = 334° ± 7°
raw map
smoothed, fitted m=1
m=0 R² = 0.26
m=1 R² = 0.39
m=2 R² = 0.31
Mathilde cell 589170196 container 511510718 σ = 12.55° κ_dir = 0.60 [0.47, 0.70] φ = 165° ± 6°
raw map
smoothed, fitted m=1
m=0 R² = 0.32
m=1 R² = 0.48
m=2 R² = 0.45

**Wendell** **cell 674570702** **container 666589599** **$\sigma = 10.18°$** **$\kappa_{dir} = 0.60$ [0.48, 0.78]** **$\varphi = 197° \pm 6°$**

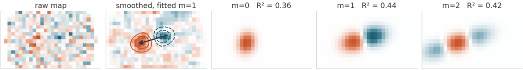


**Ting** **cell 589156275** **container 511510670** **$\sigma = 8.94°$** **$\kappa_{dir} = 0.61$ [0.56, 0.66]** **$\varphi = 199° \pm 2°$**

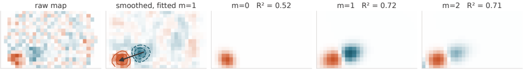


**Cornelius** **cell 662301517** **container 653125128** **$\sigma = 7.31°$** **$\kappa_{dir} = 0.65$ [0.54, 0.76]** **$\varphi = 24° \pm 7°$**

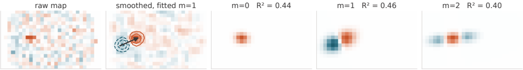


**Cassandra** **cell 517509862** **container 511510855** **$\sigma = 8.35°$** **$\kappa_{dir} = 0.67$ [0.58, 0.77]** **$\varphi = 345° \pm 5°$**

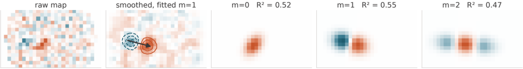


**Leontine** **cell 662279460** **container 650389885** **$\sigma = 7.65°$** **$\kappa_{dir} = 0.68$ [0.59, 0.78]** **$\varphi = 6° \pm 6°$**

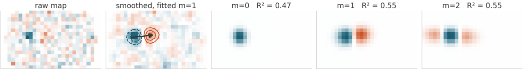


**Isadora** **cell 701720833** **container 688678764** **$\sigma = 7.93°$** **$\kappa_{dir} = 0.70$ [0.61, 0.89]** **$\varphi = 182° \pm 7°$**

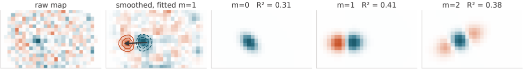

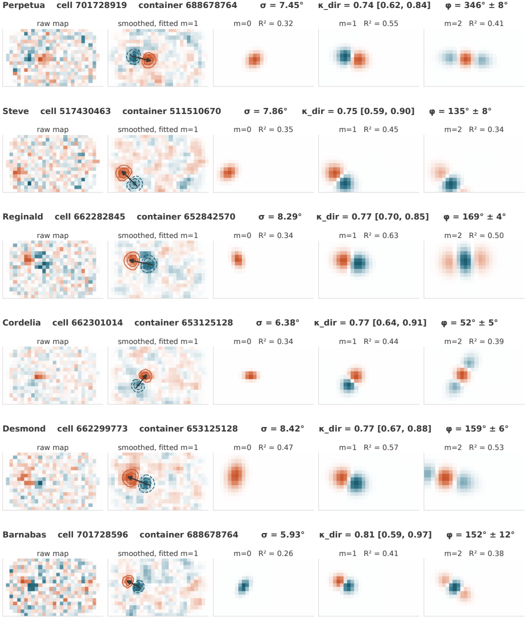
Perpetua cell 701728919 container 688678764 σ = 7.45° κ_dir = 0.74 [0.62, 0.84] φ = 346° ± 8°
raw map
smoothed, fitted m=1
m=0 R² = 0.32
m=1 R² = 0.55
m=2 R² = 0.41
Steve cell 517430463 container 511510670 σ = 7.86° κ_dir = 0.75 [0.59, 0.90] φ = 135° ± 8°
raw map
smoothed, fitted m=1
m=0 R² = 0.35
m=1 R² = 0.45
m=2 R² = 0.34
Reginald cell 662282845 container 652842570 σ = 8.29° κ_dir = 0.77 [0.70, 0.85] φ = 169° ± 4°
raw map
smoothed, fitted m=1
m=0 R² = 0.34
m=1 R² = 0.63
m=2 R² = 0.50
Cordelia cell 662301014 container 653125128 σ = 6.38° κ_dir = 0.77 [0.64, 0.91] φ = 52° ± 5°
raw map
smoothed, fitted m=1
m=0 R² = 0.34
m=1 R² = 0.44
m=2 R² = 0.39
Desmond cell 662299773 container 653125128 σ = 8.42° κ_dir = 0.77 [0.67, 0.88] φ = 159° ± 6°
raw map
smoothed, fitted m=1
m=0 R² = 0.47
m=1 R² = 0.57
m=2 R² = 0.53
Barnabas cell 701728596 container 688678764 σ = 5.93° κ_dir = 0.81 [0.59, 0.97] φ = 152° ± 12°
raw map
smoothed, fitted m=1
m=0 R² = 0.26
m=1 R² = 0.41
m=2 R² = 0.38

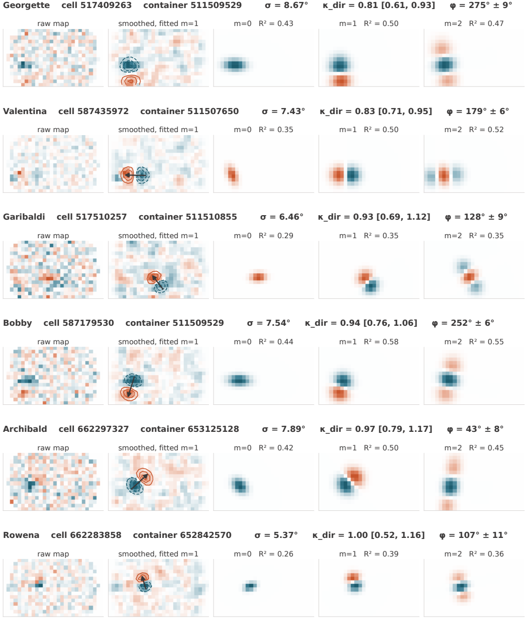
Georgette cell 517409263 container 511509529 σ = 8.67° κ_dir = 0.81 [0.61, 0.93] φ = 275° ± 9°
raw map
smoothed, fitted m=1
m=0 R² = 0.43
m=1 R² = 0.50
m=2 R² = 0.47
Valentina cell 587435972 container 511507650 σ = 7.43° κ_dir = 0.83 [0.71, 0.95] φ = 179° ± 6°
raw map
smoothed, fitted m=1
m=0 R² = 0.35
m=1 R² = 0.50
m=2 R² = 0.52
Garibaldi cell 517510257 container 511510855 σ = 6.46° κ_dir = 0.93 [0.69, 1.12] φ = 128° ± 9°
raw map
smoothed, fitted m=1
m=0 R² = 0.29
m=1 R² = 0.35
m=2 R² = 0.35
Bobby cell 587179530 container 511509529 σ = 7.54° κ_dir = 0.94 [0.76, 1.06] φ = 252° ± 6°
raw map
smoothed, fitted m=1
m=0 R² = 0.44
m=1 R² = 0.58
m=2 R² = 0.55
Archibald cell 662297327 container 653125128 σ = 7.89° κ_dir = 0.97 [0.79, 1.17] φ = 43° ± 8°
raw map
smoothed, fitted m=1
m=0 R² = 0.42
m=1 R² = 0.50
m=2 R² = 0.45
Rowena cell 662283858 container 652842570 σ = 5.37° κ_dir = 1.00 [0.52, 1.16] φ = 107° ± 11°
raw map
smoothed, fitted m=1
m=0 R² = 0.26
m=1 R² = 0.39
m=2 R² = 0.36

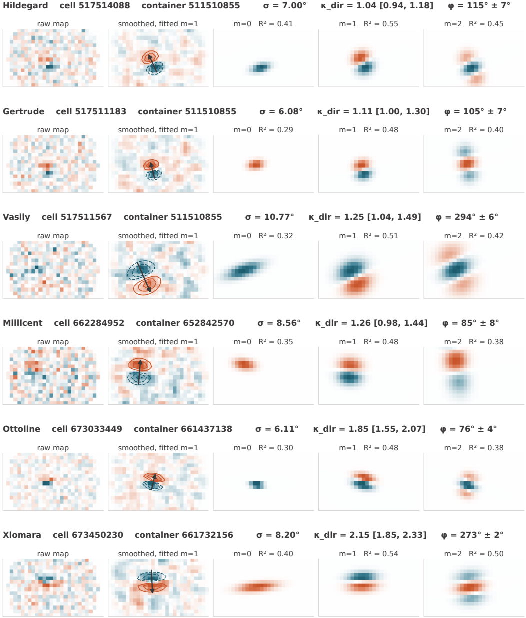
Hildegard cell 517514088 container 511510855 σ = 7.00° κ_dir = 1.04 [0.94, 1.18] φ = 115° ± 7°
raw map
smoothed, fitted m=1
m=0 R² = 0.41
m=1 R² = 0.55
m=2 R² = 0.45
Gertrude cell 517511183 container 511510855 σ = 6.08° κ_dir = 1.11 [1.00, 1.30] φ = 105° ± 7°
raw map
smoothed, fitted m=1
m=0 R² = 0.29
m=1 R² = 0.48
m=2 R² = 0.40
Vasily cell 517511567 container 511510855 σ = 10.77° κ_dir = 1.25 [1.04, 1.49] φ = 294° ± 6°
raw map
smoothed, fitted m=1
m=0 R² = 0.32
m=1 R² = 0.51
m=2 R² = 0.42
Millicent cell 662284952 container 652842570 σ = 8.56° κ_dir = 1.26 [0.98, 1.44] φ = 85° ± 8°
raw map
smoothed, fitted m=1
m=0 R² = 0.35
m=1 R² = 0.48
m=2 R² = 0.38
Ottoline cell 673033449 container 661437138 σ = 6.11° κ_dir = 1.85 [1.55, 2.07] φ = 76° ± 4°
raw map
smoothed, fitted m=1
m=0 R² = 0.30
m=1 R² = 0.48
m=2 R² = 0.38
Xiomara cell 673450230 container 661732156 σ = 8.20° κ_dir = 2.15 [1.85, 2.33] φ = 273° ± 2°
raw map
smoothed, fitted m=1
m=0 R² = 0.40
m=1 R² = 0.54
m=2 R² = 0.50
