## Supplementary figures and images for "Variabilities in the shapes of the receptive fields of simple cells in mice"

### S2 Fig

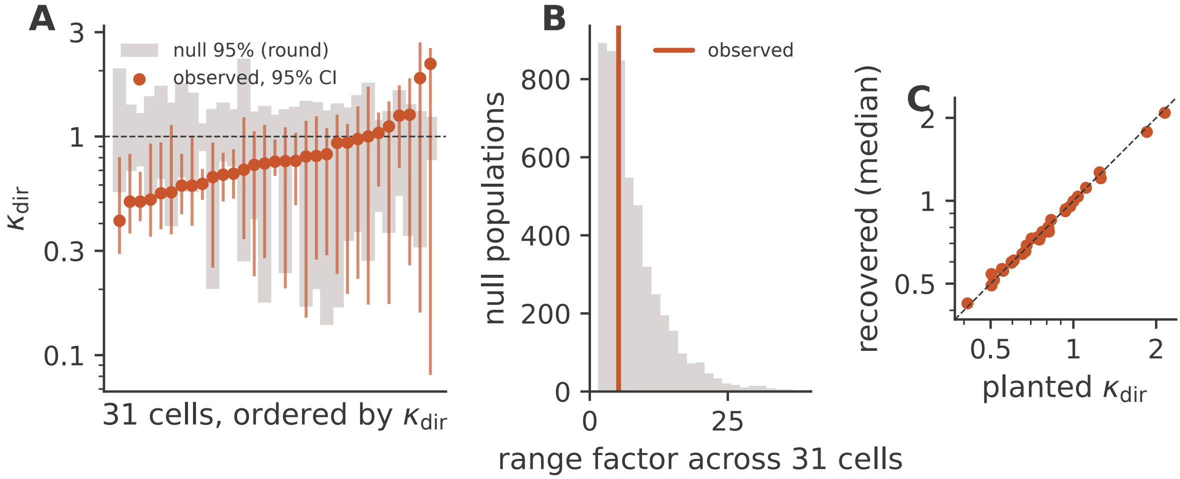
