## Supplementary material for "Variabilities in the shapes of the receptive fields of simple cells in mice": S3 Table

| cell | cell_id | container | sigma_deg | kappa_dir | kappa_dir_68_lo | kappa_dir_68_hi | kappa_dir_95_lo | kappa_dir_95_hi | phi_deg | phi_68_halfwidth_deg | r2 | p_vs_round | p_vs_round_BH |
|---|---|---|---|---|---|---|---|---|---|---|---|---|---|
| Winifred | 696526688 | 680156909 | 7.366 | 0.412 | 0.356 | 0.519 | 0.29 | 0.805 | 0 | 3.9 | 0.404 | 0.04 | 0.153 |
| Lysander | 701719883 | 688678764 | 8.637 | 0.504 | 0.448 | 0.654 | 0.36 | 0.833 | 199 | 5.4 | 0.459 | 0.01 | 0.153 |
| Bertram | 662284816 | 650389885 | 7.353 | 0.504 | 0.446 | 0.562 | 0.41 | 0.69 | 1 | 3.1 | 0.526 | 0.02 | 0.153 |
| Crispin | 589170614 | 511510718 | 7.325 | 0.515 | 0.428 | 0.669 | 0.348 | 0.929 | 204 | 6.7 | 0.385 | 0.05 | 0.153 |
| Marina | 662299371 | 653125128 | 8.552 | 0.55 | 0.454 | 0.688 | 0.376 | 0.939 | 358 | 6.4 | 0.391 | 0.04 | 0.153 |
| Fletcher | 673449540 | 661732156 | 6.309 | 0.556 | 0.485 | 0.729 | 0.357 | 1.129 | 334 | 6.5 | 0.385 | 0.069 | 0.153 |
| Mathilde | 589170196 | 511510718 | 12.548 | 0.596 | 0.469 | 0.697 | 0.441 | 0.834 | 165 | 5.6 | 0.481 | 0.059 | 0.153 |
| Wendell | 674570702 | 666589599 | 10.176 | 0.596 | 0.481 | 0.778 | 0.39 | 0.991 | 197 | 6.2 | 0.438 | 0.069 | 0.153 |
| Ting | 589156275 | 511510670 | 8.94 | 0.607 | 0.559 | 0.663 | 0.514 | 0.71 | 199 | 2.4 | 0.723 | 0.02 | 0.153 |
| Cornelius | 662301517 | 653125128 | 7.308 | 0.652 | 0.539 | 0.761 | 0.251 | 0.939 | 24 | 7.0 | 0.464 | 0.05 | 0.153 |
| Cassandra | 517509862 | 511510855 | 8.35 | 0.669 | 0.583 | 0.766 | 0.505 | 0.842 | 345 | 5.4 | 0.554 | 0.01 | 0.153 |
| Leontine | 662279460 | 650389885 | 7.653 | 0.676 | 0.592 | 0.783 | 0.519 | 0.874 | 6 | 5.7 | 0.551 | 0.03 | 0.153 |
| Isadora | 701720833 | 688678764 | 7.934 | 0.705 | 0.614 | 0.886 | 0.339 | 1.225 | 182 | 7.1 | 0.413 | 0.139 | 0.239 |
| Perpetua | 701728919 | 688678764 | 7.445 | 0.742 | 0.621 | 0.843 | 0.229 | 1.058 | 346 | 8.2 | 0.547 | 0.089 | 0.162 |
| Steve | 517430463 | 511510670 | 7.857 | 0.752 | 0.593 | 0.9 | 0.278 | 1.13 | 135 | 8.2 | 0.452 | 0.248 | 0.365 |
| Reginald | 662282845 | 652842570 | 8.293 | 0.767 | 0.7 | 0.847 | 0.66 | 0.97 | 169 | 4.4 | 0.632 | 0.059 | 0.153 |
| Cordelia | 662301014 | 653125128 | 6.384 | 0.772 | 0.639 | 0.911 | 0.202 | 1.101 | 52 | 5.4 | 0.445 | 0.178 | 0.291 |
| Desmond | 662299773 | 653125128 | 8.417 | 0.774 | 0.674 | 0.88 | 0.485 | 1.039 | 159 | 6.2 | 0.574 | 0.089 | 0.162 |
| Barnabas | 701728596 | 688678764 | 5.929 | 0.81 | 0.591 | 0.973 | 0.148 | 1.18 | 152 | 12.2 | 0.411 | 0.505 | 0.602 |
| Georgette | 517409263 | 511509529 | 8.668 | 0.815 | 0.61 | 0.927 | 0.273 | 1.238 | 275 | 9.2 | 0.495 | 0.327 | 0.44 |
| Valentina | 587435972 | 511507650 | 7.431 | 0.83 | 0.712 | 0.948 | 0.286 | 1.092 | 179 | 5.8 | 0.498 | 0.238 | 0.365 |
| Garibaldi | 517510257 | 511510855 | 6.459 | 0.935 | 0.69 | 1.12 | 0.235 | 1.26 | 128 | 9.3 | 0.349 | 0.683 | 0.784 |
| Bobby | 587179530 | 511509529 | 7.544 | 0.937 | 0.758 | 1.058 | 0.191 | 1.141 | 252 | 6.0 | 0.58 | 0.762 | 0.826 |
| Archibald | 662297327 | 653125128 | 7.895 | 0.973 | 0.794 | 1.173 | 0.223 | 1.374 | 43 | 7.6 | 0.505 | 0.851 | 0.88 |
| Rowena | 662283858 | 652842570 | 5.369 | 1.001 | 0.524 | 1.156 | 0.17 | 1.692 | 107 | 11.2 | 0.391 | 0.98 | 0.98 |
| Hildegard | 517514088 | 511510855 | 6.998 | 1.038 | 0.936 | 1.178 | 0.59 | 1.288 | 115 | 6.6 | 0.553 | 0.772 | 0.826 |
| Gertrude | 517511183 | 511510855 | 6.083 | 1.112 | 0.996 | 1.302 | 0.172 | 1.447 | 105 | 7.0 | 0.482 | 0.307 | 0.432 |
| Vasily | 517511567 | 511510855 | 10.768 | 1.246 | 1.035 | 1.492 | 0.718 | 1.715 | 294 | 6.3 | 0.507 | 0.366 | 0.473 |
| Millicent | 662284952 | 652842570 | 8.563 | 1.257 | 0.977 | 1.442 | 0.258 | 1.845 | 85 | 8.1 | 0.477 | 0.396 | 0.491 |
| Ottoline | 673033449 | 661437138 | 6.107 | 1.85 | 1.55 | 2.072 | 0.157 | 2.694 | 76 | 3.9 | 0.475 | 0.089 | 0.162 |
| Xiomara | 673450230 | 661732156 | 8.203 | 2.149 | 1.849 | 2.326 | 0.081 | 2.538 | 273 | 2.3 | 0.535 | 0.03 | 0.153 |
