## Supplementary material for "Variabilities in the shapes of the receptive fields of simple cells in mice": S4 Text

### S4 Text. Scale–elongation coupling across the full order-corrected population.

Scale and elongation are predicted by the theory to vary independently, as two separate degrees of freedom, so any association between them is a departure from the theory rather than a confirmation of it. The main text reports the result for the 31 verified first-order simple cells, among which no significant association is found. For completeness we give here the corresponding analysis across the full order-corrected population ($n$ = 1,017), together with the control for the pre-smoothing that motivates our decision not to interpret it as a property of the simple cells.

#### Population-level association

Across the full order-corrected population, scale and elongation co-varied positively (Pearson $r = 0.52$, $p < 10^{-70}$; Spearman $\rho = 0.54$, $p < 10^{-70}$; $n$ = 1,017; S4 Fig). This population, however, pools neurons of different derivative orders and types, the large majority of which are zero-order ($m$=0) fits rather than verified simple cells, so an association measured across it cannot be attributed to the simple-cell family.

#### The pre-smoothing generates part of the coupling

The same pre-smoothing that compresses elongation can also generate a coupling of this kind from nothing, so we tested that before interpreting it. Smoothing adds the same quantity to both axes of every cell, which is a large share of a small receptive field and a small share of a large one; small receptive fields are therefore recovered rounder than they are while large receptive fields are recovered nearly correctly, and a population in which every cell had the same elongation would still show scale and elongation rising together. We repeated the correlation with that contribution removed from both axes. Among the receptive fields well clear of the smoothing scale, the coupling survives but is reduced: for the 452 cells whose minor axis exceeds $1.5\,\sigma_{\mathrm{smooth}}$ it is $r = +0.41$ corrected against $r = +0.54$ uncorrected, and for the 177 cells beyond $2\,\sigma_{\mathrm{smooth}}$, $r = +0.50$ against $r = +0.56$. Including cells at the floor weakens it substantially ($r = +0.16$ over all 989 cells with usable axis estimates), but that is the correction failing rather than the coupling: for a cell whose minor axis is 3.66°, just above $\sigma_{\mathrm{smooth}} = 3.49°$, the corrected minor axis is 1.11° and the corrected ratio is 7.5, and shifting the measured axis by 0.1° moves that ratio to 5.9. Near the floor the correction subtracts two almost equal quantities, so what remains is largely measurement error. The coupling is therefore present in the range where it can be measured, while its magnitude depends on where that range is cut and should not be read as an estimate.

#### Interpretation

What a residual coupling would mean, if it holds, is an open question. One possibility is that the smallest scale a population represents is set by the acuity available at each retinotopic location, so that the finest scale level and the shapes built on it are chosen together rather than independently. We found no support for that here, since the receptive field position within the stimulus display predicted neither scale ($r = 0.04$) nor elongation ($r = 0.08$) to any useful degree, but display position is a poor stand-in for retinotopic eccentricity and the question deserves a proper retinotopic test. Taken together with the absence of a detectable association among the verified simple cells, we treat the population-level coupling as attributable in part to the pooling of different cell types and in part to the isotropic pre-smoothing, and do not interpret it as a property of the simple-cell receptive field family.

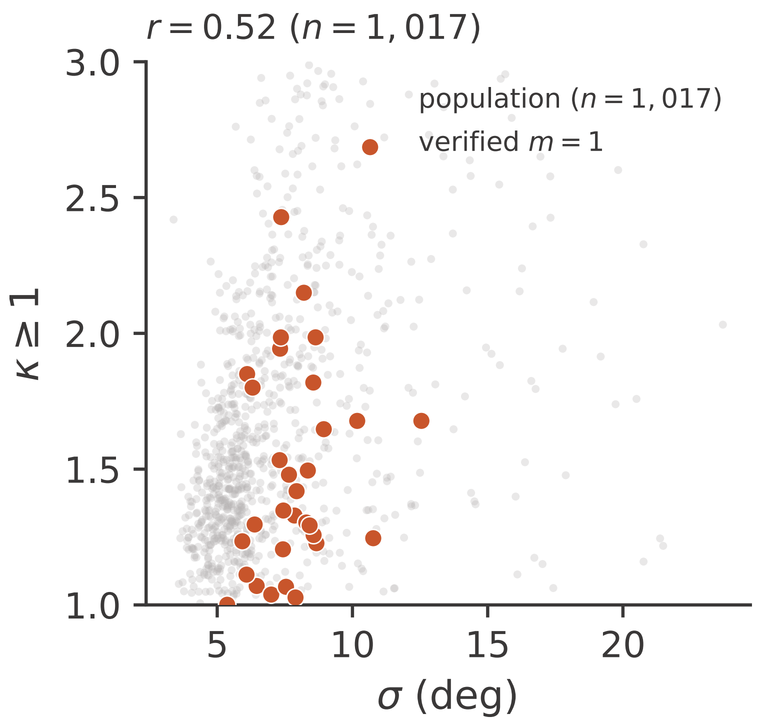


**S4 Fig 1. Scale against elongation across the order-corrected population.** Scale $\sigma$ against the symmetric elongation ratio $\kappa$ (defined for all derivative orders) across the order-corrected population ($n =$ 1,017), with the verified $m$=1 simple cells highlighted. The correlation $r = 0.52$ is computed on all 1,017 cells; the $\kappa$ axis is truncated at 3 for display only, and a small number of population cells with a near-floor minor axis extend higher.
